# CReasPy-cloning: a method for simultaneous cloning and engineering of megabase-sized genomes in yeast using the CRISPR-Cas9 system

**DOI:** 10.1101/647750

**Authors:** Estelle Ruiz, Vincent Talenton, Marie-Pierre Dubrana, Gabrielle Guesdon, Maria Lluch-Senar, Franck Salin, Pascal Sirand-Pugnet, Yonathan Arfi, Carole Lartigue

**Author notes:** **Corresponding author:** INRA, Centre de Recherche de Bordeaux, UMR 1332 Biologie du Fruit et Pathologie, 71, avenue Edouard Bourlaux, CS 20032, 33882 Villenave d’Ornon Cedex, France.

## Abstract

Over the last decade a new strategy was developed to bypass the difficulties to genetically engineer some microbial species by transferring (or “cloning”) their genome into another organism that is amenable to efficient genetic modifications and therefore acts as a living workbench. As such, the yeast *Saccharomyces cerevisiae* has been used to clone and engineer genomes from viruses, bacteria and algae. The cloning step requires the insertion of yeast genetic elements within the genome of interest, in order to drive its replication and maintenance as an artificial chromosome in the host cell. Current methods used to introduce these genetic elements are still unsatisfactory, due either to their random nature (transposon) or the requirement for unique restriction sites at specific positions (TAR cloning). Here we describe the CReasPy-Cloning, a new method that combines both the ability of Cas9 to cleave DNA at a user-specified locus and the yeast’s highly efficient homologous recombination to simultaneously clone and engineer a bacterial chromosome in yeast. Using the 0.816 Mbp genome of *Mycoplasma pneumoniae* as a proof of concept, we demonstrate that our method can be used to introduce the yeast genetic element at any location in the bacterial chromosome while simultaneously deleting various genes or group of genes. We also show that CReasPy-cloning can be used to edit up to three independent genomic loci at the same time with an efficiency high enough to warrant the screening of a small (<50) number of clones, allowing for significantly shortened genome engineering cycle times.

## INTRODUCTION

Genetically engineering a living organism is a key technology, for both fundamental and applied purposes. The ability to specifically delete, replace or add a sequence to a genome can be used to study the function of a given gene or group of genes^1^, or to add new or improved functions to a cell^2^.

From early groundbreaking experiments on bacterial transformation^3^, to the most recent advances in genomic engineering using targetable nucleases ^4,5^, a wide array of tools have been developed in order to introduce and maintain exogenous genetic material in an organism or to directly edit the existing genome. However, while multiple strategies are often available for the majority of model organisms, many species of biological interest still lack efficient genetic engineering tools^6^.

Over the last decade, it was proposed that these difficulties could be bypassed by transferring a whole genome to edit it into another cell where efficient tools are available^7^. This approach was in particular developed using the yeast *Saccharomyces cerevisiae* as a genetic workbench to manipulate whole bacterial chromosomes. This process was also applied for the creation of the first bacterial cell governed by a chemically synthesized genome^8^ and the smallest engineered living organism^9^.

Given the large array of genetic tools available in yeast, including TREC^10^, TREC-IN^11^, CRISPR-Cas9^12,13^, this genome engineering strategy has generated a growing interest. To this date, a wide range of chromosomes from different organisms have been successfully cloned in yeast, originating from viruses^14,15^, bacteria^16,17^ and algae^18^.

In order to clone a bacterial chromosome in yeast, several elements must be added to it. These “yeast elements” are: a yeast origin of replication, a yeast centromere and a selection marker. All these elements are required to drive the replication and maintenance of the foreign DNA, and are usually provided as a single cassette that can be inserted into the bacterial genome using two main strategies. The first one relies on an initial transformation of the bacteria with a transposon bearing the yeast elements^7^. While selection of living transformants ensures that the random integration of the transposon has not occurred in any locus essential for *in-vitro* growth, this method cannot be used to integrate the yeast elements at a specific position of the genome. The second strategy is based on the Transformation-Associated Recombination (TAR) cloning technique^19^. The complete bacterial genome is isolated in agarose plugs, linearized by restriction, and co-transformed into yeast together with the yeast element cassette flanked by recombination arms corresponding to both sides of the restriction locus. The yeast homologous recombination mechanism then circularize the bacterial chromosome by integration of the cassette. This approach necessitates the presence of a unique restriction site in the bacterial genome, preferably in a non-essential locus, which can be hard to find and limits the number of integration sites available.

In this study, we have developed a new strategy called CReasPy-Cloning, in order to efficiently insert the yeast elements at any desired locus of a DNA fragment to be cloned in yeast. This method expands on the logic of TAR cloning, by using the CRISPR/Cas9 system to generate a double strand break at a precise site of the genome to clone. In addition, the flexibility offered by the choice of the insertion locus allows us to simultaneously perform the cloning and edition of a bacterial genome.

To demonstrate the effectiveness of the CREASPY-cloning strategy, we cloned with a high efficiency the genome of *Mycoplasma pneumoniae* M129^20^ in yeast, while simultaneously deleting a gene encoding for a virulence factor. Analysis showed that the cloned genome was intact and essentially error-free. We successfully applied the same strategy to *Mycoplasma leachii* strain PG50^21^, in order to demonstrate that genomes cloned using CREASPY-cloning are suitable for genome transplantation. We then stretched the capability of CREASPY-cloning by successfully editing two or three distinct loci simultaneously.

## RESULTS AND DISCUSSION

### Simultaneous cloning and engineering of *Mycoplasma pneumoniae* genome in yeast

We have developed a method dubbed “CReasPy-cloning”, in order to perform the simultaneous cloning and engineering of megabase-sized genomes in yeast. Figure 1A outlines the general principle of the method. First, the yeast *Saccharomyces cerevisiae* strain VL6-48N is transformed with the plasmids pCas9 and pgRNA, allowing respectively the expression of a codon-optimized version of the Cas9 nuclease from *Streptococcus pyogenes*^12^ and a chimeric guide RNA (gRNA) merging the CRISPR RNA with the trans-activating crRNA^12,13,22,23^. Then, the yeasts are co-transformed with the purified bacterial chromosome and a linear DNA cassette comprised of the CEN-HIS yeast elements^7^ (CEN: centromere; HIS: histidine auxotrophic marker), an antibiotic resistance marker (used for an eventual back transplantation experiment later on^24^) and flanked with recombination arms corresponding to the locus to edit (Figure S1). The Cas9/gRNA complex generates a double strand break at the targeted locus on the bacterial chromosome, which is then repaired by the homologous recombination machinery of the yeast using the provided template cassette. As a result, the target locus is edited, and the bacterial genome now bears the CEN-HIS elements and is carried by the yeast as a large centromeric plasmid. After the CReasPy-cloning protocol, the yeast transformants are screened using a combination of PCR analysis and chromosome size determination by PFGE, in order to check the integrity of the cloned genome.

**Figure 1.**
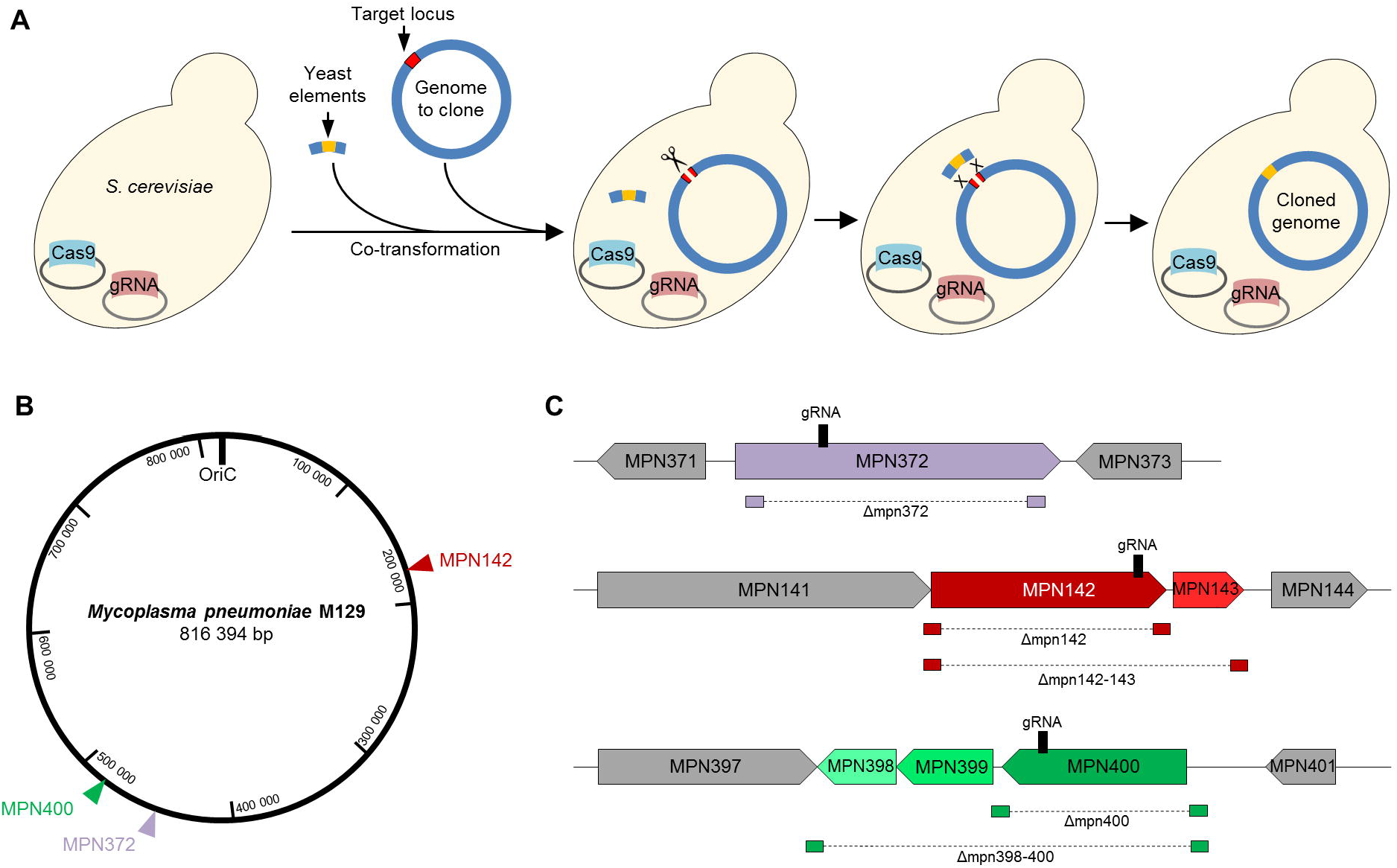
Overview of the CReasPy-cloning method and its application to *M. pneumoniae* M129 genome. (A) Schematic diagram of the experimental procedure of CReasPy-cloning. A yeast is transformed beforehand with two plasmids, allowing the expression of the Cas9 nuclease and a gRNA. This yeast is then transformed simultaneously with the genome to be cloned and a fragment of linear DNA containing the yeast elements (centromere, selection marker) flanked by two recombination arms homologous to each side of the target locus. Upon entry in the cell, the target genome is cleaved by the Cas9/gRNA complex, and subsequently repaired by the yeast homologous recombination system, using the provided linear DNA fragment as template. As a result, the bacterial genome now includes the yeast elements inserted at a precise locus (here with precise deletion of a target red gene), and is now carried by the yeast as an artificial chromosome. (B) Map of *M. pneumoniae* M129 genome. The location of the three targeted loci (MPN372, MPN142 and MPN400) is indicated by colored arrows. (C) Schematic view of the genomic regions surrounding the targeted loci. The location of the gRNA protospacer sequence is indicated by a black bar. The locations of the recombination arms are indicated by small rectangles connected by a dotted line.

As a proof of concept, we have used CReasPy-cloning to simultaneously clone and engineer the genome of the human pathogen *M. pneumoniae* M129 (816 kb), for which no efficient genome engineering tool is available. Three genes encoding virulence factors were targeted: MPN372 encoding the CARDS toxin^25^, MPN142 encoding a cytadherence protein^26,27^, and MPN400 encoding an immunoglobulin-binding protein^28^ (Figure 1B and 1C). In the case of MPN400, we targeted either the single gene or the operon it belongs to (MPN398-400). In the case of MPN142, we targeted either the single gene or part of the operon it belongs to (MPN142-143), as the other genes in the operon (MPN140-141) could be essential^29^.

Initially, a set of experiments targeting MPN372 was performed during which all the components (pgRNA, pCas9, chromosome and cassette) were co-transformed simultaneously in the yeast (Table S1). However, these experiments failed to yield positive clones. Therefore, an alternative protocol was set up, in which the pgRNA and pCas9 were transformed first, in order to give the yeast cells enough time to express the Cas9 nuclease and its partner gRNA before transformation with the bacterial chromosome. This strategy was much more efficient, and was used for the rest of the experiments described below.

First, the CReasPy-cloning was used to clone the *M. pneumoniae* genome while deleting the MPN372 gene (Figure 2). The presence of the *M. pneumoniae* genome cloned in yeast and the replacement of the target gene by the repair cassette were detected by simplex PCR. Twenty yeast transformants were screened, and all of them presented the expected profile with a single amplicon at 3,595 bp (Figure 2A). In a second step, a multiplex PCR was performed using two sets of 10 primer pairs, targeting a total of 20 loci evenly spread on the chromosome of *M. pneumoniae* (Figure S2), to check the completeness of the cloned genome. A total of 16 clones out of the 20 tested presented the expected 10 bands profile with both sets of primers (Figure 2B). Four clones (6.6, 6.7, 7.5 and 7.8) lacked one or more amplicons in one or both multiplex PCR, indicating that either one or several large parts of the bacterial genome had been lost during the CReasPy-cloning process. The cause of these large deletions in genomes that harbor the repair cassette at the correct location, is currently unknown, but might be linked the documented off-target activity of Cas9^30–32^. Indeed, if the nuclease cleaves the DNA at undesired loci, the yeast would need to perform a homologous recombination between two similar sequences elsewhere in the genome in order to maintain it. This phenomenon could be bolstered by the effective presence of multiple repeated regions in the genome of *M. pneumoniae*^33–35^. The final transformants screening was based on the analysis of large DNA fragments generated by enzymatic restriction of the edited chromosome followed by PFGE. Among the 16 yeast clones validated by multiplex PCR, six were randomly selected and checked (Figure 2C). Five clones presented the expected profile, identical to that of the non-edited genome, with two fragments at 711 kbp and 105 kbp, for a chromosome size of 816 kbp.

**Figure 2.**
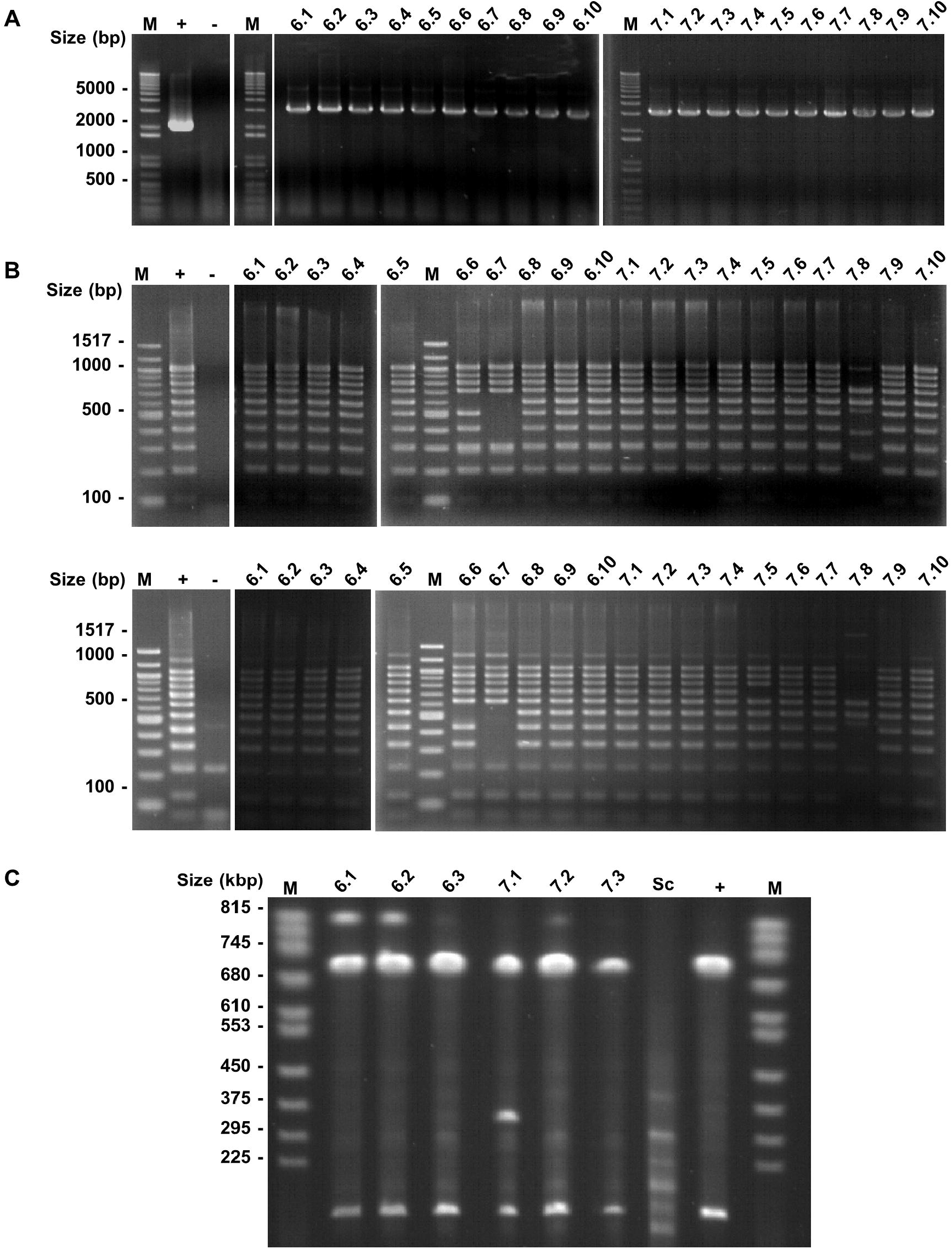
Screening of yeast transformants generated by CReasPy-cloning of *M. pneumoniae* M129 genome and deletion of MPN372. The images used to produce panels A-C are spliced from multiple gels in order to display the same 20 yeast clones. Original images are available in Supplementary Figure S3. (A) The presence of the *M. pneumoniae* genome in the yeast and the desired target replacement were checked by simplex PCR analysis. Using *M. pneumoniae* specific primers flanking the target gene, a 3,595 bp amplicon was expected in properly edited clones, whereas a fragment of 2,020 bp was expected for wild types clones. (B-C) The completeness of the *M. pneumoniae* genome cloned in yeast was assessed by multiplex PCR using two sets of primers (Top: set 1, Bottom: set 2). Each set consisted in ten pairs of primers evenly distributed around the genome of *M. pneumoniae*, allowing the simultaneous amplification of ten fragments ranging from 100 to 1,000 bp (set 1) and from 125 to 1,025 bp (set 2) in 100 bp increments. Clones carrying *M. pneumoniae* genomes without major rearrangement displayed a ten bands profile identical to the one obtained in the positive control. “M”: DNA Ladder; “+”: *M. pneumoniae* M129 gDNA; “−”: negative control without DNA. (C) The sizes of the *M. pneumoniae* genomes cloned in yeast were assessed by enzymatic restriction and Pulsed Field Gel Electrophoresis (PFGE). Digestion of the bacterial genome with the restriction enzyme NotI-HF should release two linear DNA fragments of 711 kbp and 105 kbp. “M”: PFGE DNA ladder; “Sc”: *S. cerevisiae* VL6-48N; “+”: *M. pneumoniae* M129.

In order to demonstrate the robustness of the CReasPy-cloning editing method, the same process was applied for the targeted deletions listed in Table 1. For the deletions MPN400 and MPN398-400, 100% of the 20 tested clones were found positive by simplex PCR. When we targeted the MPN142 and MPN142-143 regions, 95% and 80% of the screened clones had the expected simplex PCR profiles, respectively (Table 1). At the next step, 79%, 69%, 50% and 70% of the clones validated by simplex PCR were found positive in multiplex PCR, for the deletion of MPN142, MPN142-143, MPN400 and MPN398-400, respectively. Finally, 67%, 67%, 83% and 83% of the clones tested by PFGE were positive, for MPN142, MPN142-143, MPN400 and MPN398-400, respectively (Table 1).

Overall, the CReasPy-cloning method we developed exhibits a high efficiency, as only a small number of clones (around 20) have to be screened to identify those carrying the expected edited bacterial chromosome. Using a yeast elements cassette which does not contain an Autonomously Replicating Sequence^36^ (ARS) was identified as a key factor in reaching this high efficiency. The ARS acts as an origin of replication for the artificial yeast chromosome^7^ and must contain a copy of the essential 11 bp ARS Consensus Sequence: 5’-WTTTAYRTTTW-3’. It has been reported that removing the ARS sequence from TAR vector increased the efficiency of transformation^37^. This improvement was linked to the fact that the recombination template tends to circularize instead of integrating into the bacterial chromosome during yeast transformation. This circular element bearing the ARS sequence is efficiently replicated in yeast, leading to false-positive clones. In our case, the removal of the ARS sequence did not jeopardize the replication of the *M. pneumoniae* chromosome by the yeast machinery, as the 11 bp ARS Consensus Sequence is present 19 times in the bacterial genome. During an initial experiment, we performed CReasPy-cloning experiment with an ARS-containing or an ARS-less recombination template (Table S2), and obtained better results with the latter.

In addition, several other genome cloning experiments were also performed in the yeast *S. cerevisiae* strain W303a, as this strain and the strain VL6-48N were both used in previous studies focused on the cloning and edition of mycoplasma genomes in yeast^7,38,39^. Overall, similar transformation efficiencies were obtained for both strains (data not shown). Strain VL6-48N was retained for the rest of the experiments.

### Application of CReasPy-cloning to multiple *Mycoplasma* species

Following the successful development of our method for cloning and editing *M. pneumoniae* genome, we applied it on two other mycoplasma species: *Mycoplasma leachii* strain PG50 and *Mycoplasma mycoides* subsp. *mycoides* strain Afadé (*Mmm*). The aims of these experiments were to *i*) validate the CReasPy-cloning method for larger chromosomes, *ii*) re-clone mycoplasma genomes for which the yeast elements were previously inserted randomly by transposons^39^ and, *iii*) check the ability of the CReasPy-cloned genomes to be efficiently transplanted.

We succeeded in cloning *M. leachii* genome (~1 Mb) while inactivating MSB_0138, a gene encoding a beta-lactamase. We also successfully performed the cloning of *Mmm* genome (~1.2 Mb), while deleting of TS60_0301-0299, a trio of genes encoding an α-glycerolphosphate oxidase, a glycerol kinase and a glycerol facilitator. Positive yeast clones carrying edited whole genomes were obtained for both species (Table 2 and Figure S4-S5). We observed that the cloning efficiencies of *Mmm* and *M. leachii* genomes (Table 2) were in the same range as those measured for *M. pneumoniae*, indicating that the CReasPy-cloning method is well suited for chromosomes in the megabase range.

In the case of *M. leachii*, the genome CReasPy-cloned in yeast of clones 9, 34, 36, 37 and 39 were isolated and transplanted in the recipient cell *Mycoplasma capricolum*^38^, yielding between 10 and 50 colonies depending on the clone (cl. 8 yielded no transplant). Forty putative bacterial transplants were analyzed by simplex PCR (Figure S4): 36 were identified as edited *M. leachii* cells while the other 4 were identified as *M. capricolum* tetracycline spontaneous resistants^39^. These experiments confirmed the robustness of the CReasPy-cloning as a fast and efficient method for the production of mutant strains of bacteria, provided that a transplantation protocol exists for these species. As such protocols are not currently available for *M. pneumoniae* and *Mmm*, transplantation of their CReasPy-cloned genomes were not attempted during this study.

It is currently unknown whether the size of the target chromosome is a limiting factor for the “in-yeast cloning” process. Currently, the largest genomes carried in yeast in a single piece are the 1.8 Mbp genome of *Haemophilus influenzae*^17^ and the 1.8 Mbp genome of *Spiroplasma citri*^39^. Interestingly, a recent experiment of chromosome fusion in *S. cerevisiae* has shown that the yeast can readily replicate a single chromosome of 11.8 Mbp^40^. This result suggests that the limit might not be the size of the chromosome to clone, but rather how much total DNA the yeast can carry (6.8% excess for *M. pneumoniae*, 15% excess for *H. influenzae*). The advent of synthetic strains of *S. cerevisiae* could offer some insights, as these cells have significantly reduced genomes (around 1.1 Mbp removed from Sc2.0)^41^ and might allow to clone larger chromosomes.

### Simultaneous cloning and multi-target edition of *Mycoplasma pneumoniae* genome

In order to improve our ability to efficiently create highly edited bacterial genomes, we evaluated if CReasPy-cloning could be used to perform simultaneous multi-target editions. To do so, we performed the deletion of either two or three loci: MPN372*/MPN400; MPN142-143*/MPN398-400; MPN372*/MPN142-143; MPN372*/MPN142/MPN400; MPN372*/MPN142-143/MPN398-400. For each experiment, one locus was replaced by the yeast element cassette (indicated by a “*” next to the locus name) as the other locus or loci was simply deleted by homologous recombination with a DNA template.

The multi-target protocol is based on the same principle as the single-target CReasPy-cloning, with few modifications (Figure 3). First, the yeasts are “primed” by transformation with pCas9 and a multi-guide pgRNA. To reduce the possibility of off-target cleavage by the Cas9 nuclease, we elected to use the mutant eSpCas9 which was shown to have a higher specificity^42^ than the wild-type. Afterwards, the cells are co-transformed with *M. pneumoniae* chromosome and 2 or 3 recombination templates. One of the template contains the CEN-HIS yeast elements, while the other or other two are clean deletion cassettes comprised of two juxtaposed 500 bp regions identical to the loci surrounding the gene to delete. These cassettes are tedious to produce (here by overlap PCR), but the large size of their recombination arms have a positive impact on the recombination efficiency^37^, which offset the lack of a selection marker. Alternatively, 2×45 bp deletion cassettes, easily produced by annealing of two 90 bp oligonucleotides, can be used and have been shown to be efficient to engineer the genome of *S. cerevisiae*^12^ or to edit a mycoplasma genome cloned in *S. cerevisiae*^13^. After the multi-target CReasPy-cloning, yeast transformants were screened using the same process as in the single-target protocol.

**Figure 3.**
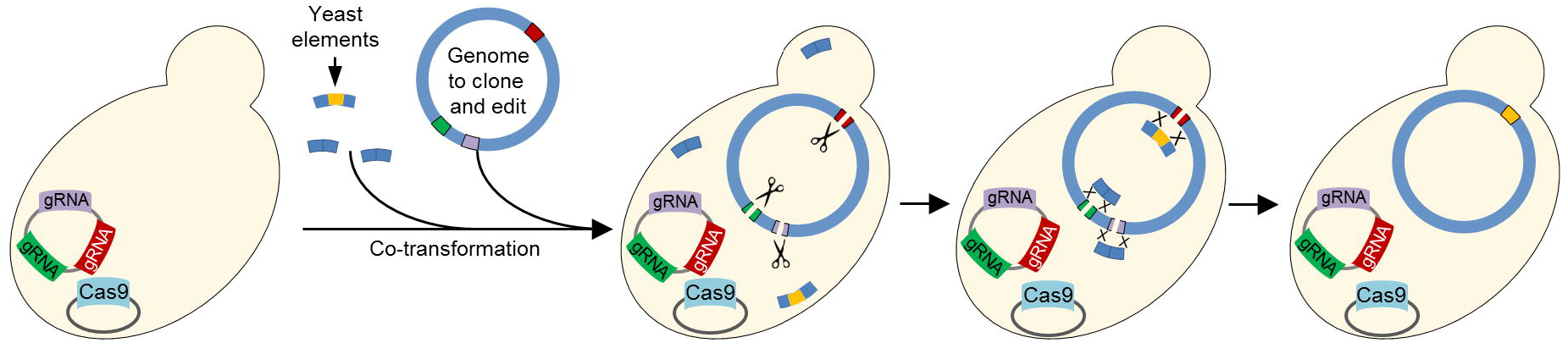
Overview of the multi-target CReasPy-cloning method. Yeast cells are transformed beforehand with two plasmids, allowing the expression of the Cas9 nuclease and multiple gRNA. The yeast is then transformed simultaneously with the target genome and three linear DNA fragments. One fragment contains the yeast elements (centromere, selection marker) flanked by two regions homologous to the target locus. The two remaining fragments are each comprised of two regions homologous to the target locus concatenated. Upon entry in the cell, the heterologous genome is cleaved at multiple loci by the Cas9/gRNA complexes, and subsequently repaired by the yeast homologous recombination system using the provided linear DNA templates. As a result, the heterologous chromosome now bears the yeast elements, inserted at a precise locus, and is now carried by the yeast as an artificial chromosome. In addition, the two other loci are edited, according to the experimenters design.

Figure 4 illustrates an example of screening for the triple deletion MPN372*/MPN142/MPN400. Twenty yeast clones were first checked by simplex PCR for the deletion of MPN372 and its replacement by the yeast elements cassette (Figure 4A), which yielded 14 positive clones with the expected amplicon at 3,596 bp. These positive clones were subsequently screened by PCR for the deletion of MPN142 (Figure 4B), yielding four positive clones presenting the expected 965 bp band. Finally, the deletion of MPN400 was assessed (Figure 4C), resulting in three positive clones presenting the expected 2,156 bp band. The three clones obtained for the triple deletion were successfully validated by multiplex PCR (Figure 4D) and PFGE (Figure 4E).

**Figure 4.**
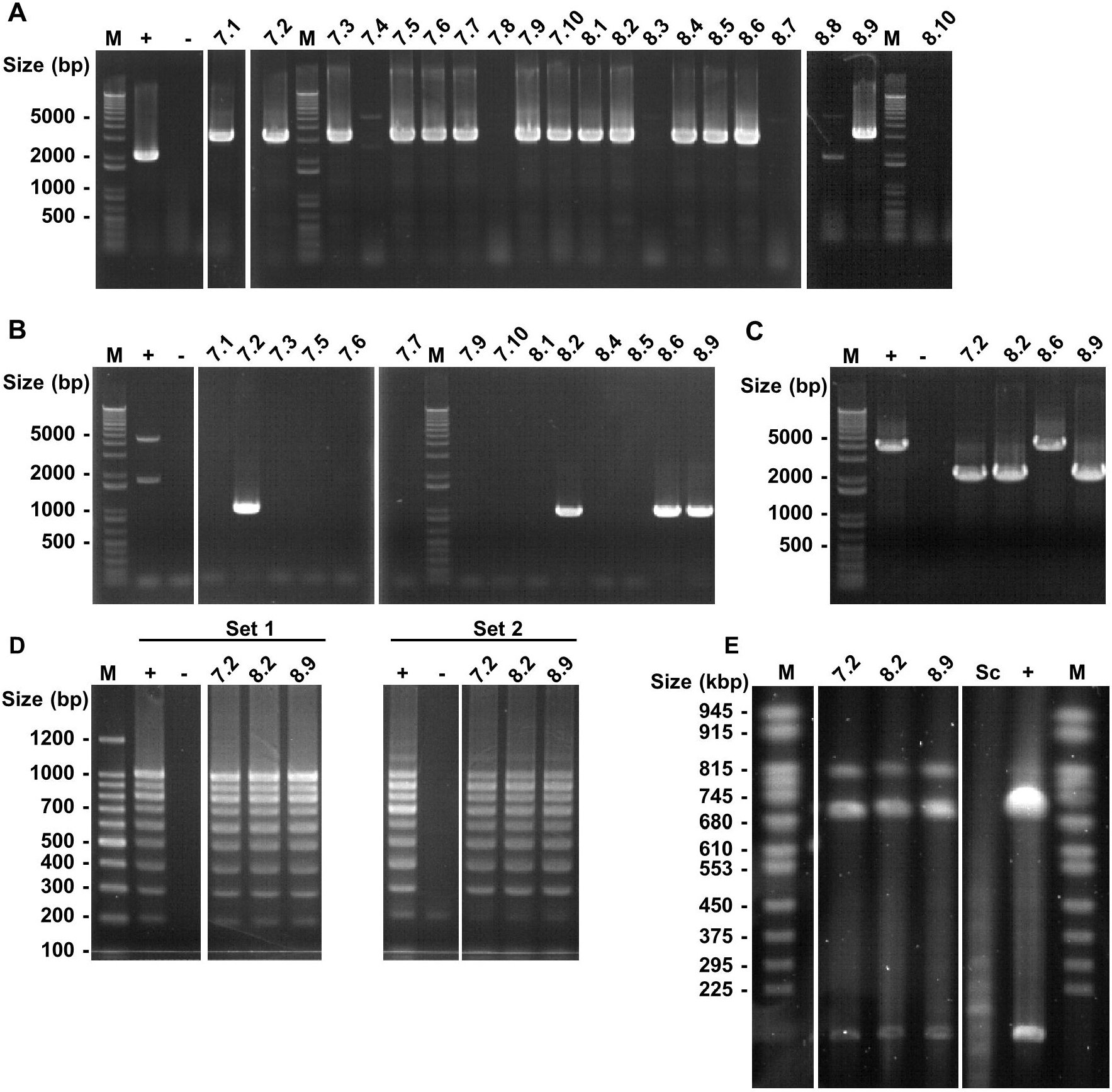
Screening of yeast transformants generated by CreasPy-cloning carrying *Mycoplasma pneumoniae* M129 genome and deletion of MPN372*/MPN142/MPN400. The images used to produce panels A-E are spliced from multiple gels in order to display the same 20 yeast clones. Original images are available in Supplementary Figure S6. (A) The presence of the *M. pneumoniae* genome in the yeast and the correct replacement of MPN372 were checked by simplex PCR analysis. Using *M. pneumoniae* specific primers flanking the target gene, an amplicon of 3,595 bp was expected in properly edited clones, whereas a fragment of 2,020 bp was expected for wild types clones. (B) The correct deletion of MPN142 was assessed by simplex PCR analysis. The expected amplicon size in clones bearing the deletion was 965 bp, whereas an amplicon of 4,598 bp was expected for clones bearing the wild-type genome. (C) The correct deletion of MPN400 was assessed by simplex PCR analysis. The expected amplicon size in clones bearing the deletion was 2,156 bp, whereas an amplicon of 4,048 bp was expected for clones bearing the wild-type genome. (D) The completeness of *M. pneumoniae* genome in yeast was assessed by multiplex PCR using two sets of primers (see Figure 2 legend). Clones carrying *M. pneumoniae* genomes without major rearrangement displayed a ten bands profile identical to the one obtained in the positive control. “M”: DNA Ladder; “+”: *M. pneumoniae* M129 gDNA; “-”: negative control without DNA. (E) The size of the *M. pneumoniae* genome in yeast was assessed by enzymatic restriction and pulsed-field gel electrophoresis (PFGE). Digestion by the restriction enzyme NotI-HF should yield two linear DNA fragments of 711 kbp and 105 kbp. “M”: PFGE DNA ladder, “Sc”: *S. cerevisiae* VL6-48N; “+”: *M. pneumoniae* M129.

All the multi-target deletions tested yielded positive clones (Table 3). As expected, due to the large number of co-transformed DNA fragments, the efficiency of the multi-target approach is significantly lower than that of the single-target CReasPy-cloning. Nonetheless, this efficiency is still high enough (minimum of 5% of the screened clones are fully validated) to warrant the screening of a relatively small set of transformants (*i.e.* <50). It is interesting to note that the majority of the negative simplex PCR showed no amplification instead of amplification of the wild-type locus, suggesting that the targeted region is absent of the final chromosome. This high rate of unwanted deletions could be caused by the stochastic nature of the transformation process. Indeed, for the CReasPy-cloning process to occur properly, each cell must receive at least one copy of each recombination template. As there is a low probability of this perfect-case scenario to occur, many cells will lack one or more templates. As a result, the transformed chromosome will be cleaved at the desired site, but in the absence of a template the recombination will happen in a spurious manner between similar loci. This suggests that, although possible in theory, stretching the CReasPy-cloning strategy to four or five targets might not be practical. Nonetheless, the ability to simultaneously clone a megabase-sized genome and edit three individual loci is still a significant improvement over existing strategies.

### Whole genome sequencing of genomes cloned in yeast by CReasPy-cloning

Whole genome sequencing was performed on one of the genome cloned and edited using CReasPy-cloning, in order to check whether our protocol had any mutagenic effects. We selected one clone from the double target experiment (MPN372*/MPN142-143 clone 4.8). Genomic DNA (including the cloned bacterial chromosome) was extracted from the yeast transformants and sequenced using both short reads (Illumina) and long reads technologies (Oxford Nanopore). As a control, the original *M. pneumoniae* M129 genome used for CReasPy-cloning experiments was re-sequenced using Illumina technology. *De novo* assembly of the genome of clone 4.8 was achieved using long and short reads, yielding a single contig. Global alignment of the clone 4.8 assembly with the expected genome design confirmed that no large deletion or chromosomal shuffling had occurred. In order to identify potential SNP or short indels, Illumina short reads from clone 4.8 and control *M. pneumoniae* M129 were mapped on the published genome sequence of *M. pneumoniae* strain M129 (Genbank ID NC_000912.1). The control clone of *M. pneumoniae* M129 used in our study presented 105 SNP and 19 indels compared to the M129 published genome. Those mutations were also identified in the genome of clone 4.8, suggesting that they occurred before the CReasPy-cloning process. Two additional mutations that were found only in clone 4.8. The first is an AAG to AAU transversion at position 182216 of the genome, leading to a K to N amino acid change in MPN141 (adhesin P1). The second is an insertion of ATGTTTG at position 452508 of the genome, in a region predicted to encode the ncRNA MPNnc041. The extent of the mutations observed in clone 4.8 suggest that the CReasPy-cloning method has no large mutagenic effect. However, SNP were expected as a result of natural genetic drift during the passages of either the mycoplasma or yeast cells in the laboratory.

### *In vitro* CReasPy-Cloning

In order to improve the flexibility of the CReasPy-cloning process, we attempted to perform the chromosome cleavage step *in vitro* (Figure S7). Indeed, this alternative strategy could offer several benefits compared to the *in vivo* method described above: *i*) it would not be necessary to “prime” the yeast with the pgRNA and pCas9 plasmids, reducing the number of transformation steps to one, *ii*) the selection markers used to maintain the pgRNA and pCas9 plasmids could be allocated for other purposes, *iii*) a large number of different gRNA could be used simultaneously without the need to build a complex multi-gRNA plasmid, *iv*) the host cell genome could not be damaged by off-target activity, as the nuclease is not present in the cell. To do so, agarose plugs containing *M. pneumoniae* chromosomes were incubated in a mix of recombinant Cas9 from *S. pyogenes* and gRNA expressed by *in vitro* transcription from oligonucleotides. The Cas9-gRNA complex is able to diffuse through the agarose matrix and cleave the chromosome at the target site. The cut chromosome was subsequently co-transformed in yeast together with a recombination cassette bearing the yeast elements.

We validated this *in vitro* CReasPy-cloning strategy by targeting MPN400, using the same recombination cassette and gRNA spacer as those used in the *in vivo* approach. Ten yeast transformants were subsequently screened as described above (Figure S8): all the checked clones were positive by simplex PCR, and four were positive by multiplex. Of these, three were validated by PFGE.

These results confirm the functionality of the *in vitro* approach, albeit with a slightly reduced efficiency compared to the *in vivo* strategy. Similar strategies, based on *in vitro* cleavage of DNA by CRISPR/Cas9 followed by capture in yeast have been highlighted by other groups^43,44^ but these approaches are currently limited to sizes below 150 kbp and are not suitable to work on whole bacterial genomes.To go on further, optimization of multiple parameters could be considered, including the concentration of Cas9 and gRNA, the incubation time and the amount of target chromosomes per agarose plug.

## CONCLUSION

In this study, we developed the CReasPy-cloning tool, to efficiently clone megabase-sized DNA molecules in *S. cerevisiae*, while simultaneously editing up to three independent loci. This approach was originally conceived in order to improve the cloning process of bacterial genomes, in particular of the genus *Mycoplasma*, and to significantly shorten the time necessary to produce highly engineered strains for academic or applied purposes^45^.

However, our method is not limited to this type of application, as its reliance on the CRISPR/Cas9 system and homologous recombination makes it highly versatile and able to process DNA from different origins and of various sizes, as the entire genomes of some viruses, bacteria, chloroplasts and mitochondria or parts of more complex genomes as those of eukaryotic cells.

In particular, we propose that CReasPy-cloning could be a valuable tool to capture intact chromosomes of uncultivable microorganisms from environmental samples (Figure S10). A target sequence could be selected based on previous or co-occurring metagenomics analysis, even if only small contigs are available. By producing an appropriate gRNA and recombination cassette, the chromosome of interest could in effect be “fished-out” by the yeast from the complex mixture of environmental DNA. Once cloned in yeast, these genomes could be easily sequenced and assembled in a single contig, which might not be possible using shotgun metagenome sequencing for rare organisms^46^.

## METHODS

### Yeast and bacterial strains, culture conditions

*Saccharomyces cerevisiae* strain VL6-48N (*MAT***a**, *his3-*Δ*200*, *trp1-*Δ*1*, *ura3–52*, *lys2*, *ade2– 101*, *met14)* is grown at 30°C in YPDA medium (Clontech). Yeast transformants are selected by growth in Synthetic Defined (SD) medium depleted for one or several amino-acids: SD-Trp, SD-His or SD-Trp-Ura (Clontech). *Mycoplasma pneumoniae* strain M129 (ATCC 29342), *Mycoplasma leachii* strain PG50 (*M. leachii*) and *Mycoplasma mycoides* subsp. *mycoides* strain Afadé (*Mmm*) are grown at 37°C in SP5 medium^39^. *Escherichia coli* strains DH10B, strain NEB5-α or strain NEB10-β used for plasmid cloning are grown at 37°C in lysogenic broth (LB) medium supplemented with 100 µg.mL^−1^ of ampicillin.

### Oligonucleotides

All the oligonucleotides used for this study are supplied by Eurogentec and are described in Table S3.

### Construction of gRNA plasmids for simple target deletion

The gRNA targeting the loci MPN372, MPN142 and MPN400 in the *M. pneumoniae* genome, MSB_A0138 in the *M. leachii* genome and TS60_0301-0299 in the *Mmm* genome, are designed using the SSC tool (http://crispr.dfci.harvard.edu/SSC/) with default parameters. The corresponding pgRNA plasmids are constructed following the protocol described in Tsarmpopoulos *et al*, 2015^13^. Briefly, the plasmid p426-SNR52p-gRNA.AarI.Y-SUP4t ("pgRNA") contains all the elements necessary for the expression of the gRNA in yeast^12^. The spacer component of the gRNA can be swapped out by restriction of the plasmid using AarI, followed by ligation of annealed oligonucleotides pairs. The resulting plasmids are transformed in *E. coli* and sequence verified.

### Construction of gRNA plasmid for triple targets deletion

The cassettes allowing the expression of the gRNAMPN142 and gRNAMPN400 are first amplified by PCR from the plasmids pgRNAMPN142 and pgRNAMPN400 respectively. These two cassettes are subsequently cloned in the plasmid pgRNAMPN372, using the Gibson Assembly Cloning Kit (NEB). The resulting plasmid pgRNA-triple-target is transformed in *E. coli* and sequence verified.

### Construction of gRNA plasmids for double targets deletion

The cassettes allowing the expression of the gRNAMPN142 and the gRNAMPN400 are PCR amplified from the plasmid pgRNA-triple-target. The resulting fragments are cloned in the linearized pgRNAMPN372 (described above), using the Gibson Assembly Cloning Kit (NEB), producing the plasmids pgRNAMPN372-142 and pgRNAMPN372-400 respectively. The pgRNAMPN142-400 plasmid is constructed by removing the gRNAMPN372 from the pgRNA-triple-target plasmid using the Q5 Site-Directed Mutagenesis Kit (NEB).

### Plasmid transformation in yeast

Yeast are transformed using the lithium acetate protocol optimized by Gietz *et al* (1995)^47^. One µg of purified plasmid is used for each transformation, and transformants are selected for auxotrophy complementation (gRNA plasmid: -Ura and Cas9 plasmid: -Trp).

### Construction of recombination templates

Recombination templates containing the yeast elements are produced by PCR amplification of the ARS/CEN/HIS/PSPuro or CEN/HIS/PSPuro loci from the plasmid pMT85-PRS-PSpuro (Figure S11), using the Advantage 2 Polymerase kit (Clontech). Complementary 60 bp-ends to the target sequence on *M. pneumoniae* genome are added to the extremities of the cassettes by using 5’-tailed PCR primers. Recombination templates comprised only of the regions flanking the loci to delete (without yeast element) are produced either by: (a) annealing of two 90 bp oligonucleotides, with 45 bp corresponding to each side of the region to be knocked-out (initial denaturing step of 5 min at 95°C and controlled cooling to 16°C with a ramp of 0.1°C.s^−1^), or (b) overlap PCR of two DNA fragments of 500 bp amplified from the *M. pneumoniae* genome using the Q5 High-Fidelity DNA Polymerase, with each 500 bp PCR fragments corresponding to the DNA sequence surrounding the loci to be deleted.

### Isolation of mycoplasma chromosomes in agarose plugs

Mycoplasma cells are grown in SP5 media, and harvested either by scrapping the culture flask bottom in HEPES-Sucrose buffer (HEPES 8mM, sucrose 272mM, pH 7.4) for *M. pneumoniae* or by centrifugation and resuspension in T/S buffer (10mM Tris pH 6.5, 500 mM sucrose) for *M. leachii* and *Mmm*, The cell suspension is then embedded in 1% low-melt agarose plugs and treated using the CHEF Mammalian Genomic DNA Plug Kit (Biorad), according to the manufacturer’s protocol^38,39^. This preparation method yields agarose plugs that contain the isolated and intact mycoplasma chromosomes. The quality of the genomic DNA is checked by digesting trapped genomes with 50 units of restriction enzymes from NEB per half of agarose plug (NotI-HF for *M. pneumonia*, XhoI for *M. leachii* and BssHII for *Mmm*) followed by a pulsed-field gel electrophoresis. Prior to yeast transformation, mycoplasma genomes are released by digestion of the agarose matrix with three units per plug of β-Agarase I (NEB), and the DNA concentration is measured using an Epoch™ Microplate Spectrophotometer (BioTek™).

### Yeast transformation with *Mycoplasma* chromosomes and recombination templates

Yeast cells carrying the pCas9 and pgRNA plasmids are transformed as described by Kouprina and Larionov (2008)^19^. For the simple target deletion experiments, 100 µL of yeast spheroplasts are mixed with 2 µg of genomic DNA and 300 ng of recombination template containing the yeast elements. For the multiple target deletion experiments, 100 µL of yeast spheroplasts are mixed with 2 µg of genomic DNA, 300 ng of recombination template containing the yeast elements and 0.5 or 1 µg of each recombination template without the yeast elements. After transformation, the yeast cells are selected on SD-His solid agar plates containing 1 M of sorbitol, for 4 days at 30°C. Individual colonies are picked and streaked on SD-His plates and incubated 2 days at 30°C. Then, one isolated colony per streak is patched on the same medium and incubated for 2 days at 30°C.

### Screening of yeast transformants carrying *Mycoplasma* genome

Total genomic DNA is extracted from yeast transformants according to Kouprina and Larionov (2008)^19^. Positive clones are screened for both the presence of the *Mycoplasma* genome and the correct deletion of the target gene by PCR, using the Advantage 2 Polymerase kit (Clontech) and specific primers located on either side of the target locus. Yeast transformants are then screened for bacterial genome completeness by multiplex PCR using two sets of PCR primers for *M. pneumoniae* (Table S3) and one set of primers for *M. leachii* and for *Mmm* (Table S4 and S5 respectively). Each set is comprised of ten pairs of primers evenly distributed across the bacterial genomes allowing the simultaneous amplification of ten fragments ranging from ~100 to ~1000 bp, in ~100 bp increment. Clones carrying mycoplasma genomes with no major rearrangements display a characteristic ten bands ladder-profile with each primer set. The multiplex PCR are performed using the Qiagen Multiplex PCR Kit according to the manufacturer’s instructions.

Yeast clones appearing positive by multiplex PCR are ultimately analyzed by restriction digestion and pulsed-field gel electrophoresis (PFGE) to assess the size of the mycoplasma chromosome. To do so, yeast cells are grow in SD-His media, harvested, embedded in agarose plugs and lysed by treatments with zymolyase, proteinase K and detergents to yield intact chromosomes. At this stage, yeast plugs carrying *M. pneumoniae* genomes are treated slightly differently compared to those containing *M. leachii* or *Mmm* genomes. For *M. pneumoniae*, the agarose-embedded yeast DNA is digested overnight with SgrDI (50U/ ½ plug) from Thermo Scientific™ and submitted to a first PFGE (1% agarose, 0.5X TBE) during 24h, with a switch time of 50-90 s, at 6 volts.cm^−1^, an angle of 120° and a temperature of 14°C. The plugs are then treated overnight with Plasmid-Safe™ ATP-dependent DNase (50U/ ½ plug) from Epicentre and loaded on a standard gel (1% agarose, 1X TAE, 120 min at 120 volts). The plugs are finally digested overnight with NotI-HF (50U/ ½ plug) and submitted to a second PFGE for 22h (using the parameters previously described). The two first steps, both performed to electrophorese the linear yeast DNA out of the plugs while preserving the circular *M. pneumoniae* chromosome, turned out to be necessary to visualize the *M. pneumoniae* genome on gel at the end of the process. Yeast plugs carrying *M. leachii* or *Mmm* genomes are hydrolysed with a cocktail of restriction enzymes (AsiSI, FseI and RsrII) and submitted to classical electrophoresis. Then, after the electro-removal of the yeast linear chromosomes, the DNA remaining in plugs is restricted with XhoI (*M. leachii*) and BssHII (*Mmm*) and submitted to PFGE. Pulse times are ramped from 60 to 120 s for 24 h at 6 volts.cm^−1^. Agarose gels are stained with SYBR™ Gold Nucleic Acid Gel Stain (Invitrogen™) and PFGE patterns are scanned using the Vilbert Lourmat™ E-BOX™ VX2 Complete Imaging system.

### Genome sequencing

Genomic DNA of yeast MPN372*/MPN142-143 clone 4.8 clone harboring edited *M. pneumoniae* chromosome was purified using Qiagen Genomic-Tips 100/G and Genomic DNA Buffers as described in Istace *et al* (2017)^48^. DNA sequencing was performed at the Genome Transcriptome Facility of Bordeaux (https://pgtb.cgfb.u-bordeaux.fr) on a GridION (Oxford Nanopore, release 18.02, flowcell R9.4.A RevD) sequencer and a MiSeq sequencer (Illumina) using paired ends libraries. ONT sequencing generated ~1,870,000 reads whereas ~3,000,000 read pairs were obtained with Illumina technology. De novo assembly process included the following steps: (1) filtering long reads with Filtlong v0.2.0 (sequences > 1000bp score phred >= 9, https://github.com/rrwick/Filtlong), (2) selection of long reads mapping *M. pneumoniae* genome with Minimaps 2:v2.15-r905^49^ (73676 reads selected), (3) assembly using Minimap/minimiasm^50^ (Miniasm 0.3-r179: https://github.com/lh3/miniasm), correction and polishing with Racon v1.3.1^51^ (4 cycles) and Pilon^52^ v1.23 (4 cycles) tools. Illumina short reads used for polishing were trimmed with Trimmomatic^53^ 0.38 (MINLEN:35, SLIDINGWINDOW:5:25). Whole genome alignment was performed with progressive MAUVE (version 20150226 and ref PMID:20593022). For short reads mapping onto M129 reference genome, data processing including quality check, trimming, alignment with BWA (Galaxy Version 1.2.3) and variant calling using Varscan (Galaxy Version 0.1) was completed using Galaxy instance (https://usegalaxy.org/)^54^.

### In vitro CReasPy-cloning

Agarose plugs containing M. pneumoniae chromosomes are incubated overnight at 37°C in presence of 1 µL of Cas9 Nuclease from Streptococcus pyogenes (M0386T, NEB) and 21 µg of gRNA transcripts produced by in vitro transcription of oligonucleotides (HiScribeTM T7 High Yield RNA Synthesis kit from NEB). Plugs are then treated overnight at 50°C with 10 µL of proteinase K, washed several times in Tris 20mM pH8, EDTA 50mM and released from agarose gel by treatment with β-Agarase I (3 units/plug). Yeast cells are finally transformed as previously described: 100µL of yeast spheroplasts are mixed with ~4 µg of genomic DNA (~50 µL of gDNA plus 50µL of TE 1x) and 300 ng of recombination template containing the yeast elements. The yeast transformants are selected and screened as described above.

## ABBREVIATION

CRISPR/Cas: Clustered Regularly Interspaced Short Palindromic Repeats/CRISPR-associated systems
PAM: Protospacer Adjacent Motif
PFGE: Pulse Field Gel Electrophoresis
TREC: Tandem Repeat Coupled with Endonuclease Cleavage
TREC-IN: TREC-assisted gene knock-IN
TAR cloning: Transformation-Associated Recombination cloning

## AUTHOR INFORMATION

Corresponding author: INRA, UMR 1332 de Biologie du Fruit et Pathologie, F-33140 Villenave d’Ornon, France Université de Bordeaux, UMR 1332 de Biologie du Fruit et Pathologie, F-33140 Villenave d’Ornon, France Tel: +33 5 57 12 23 59; Fax: +33 5 57 12 23 69;

## AUTHOR CONTRIBUTION

E.R., M.L-S., Y.A. and C.L. conceived and designed the research.

E.R., V.T., M-P.D., G.G and C.L. performed laboratory experiments and analyzed the *in vitro* data.

P.S.-P. performed genome sequencing design and subsequent analyses.

F. S. performed de novo assembly of mycoplasma genome.

E.R., Y.A. P.S.P. and C.L. wrote the paper.

All authors approved the final version of the paper.

## ACKNOWLEDGMENTS

The authors thank the Genome Transcriptome Facility of Bordeaux for genome sequencing (https://pgtb.cgfb.u-bordeaux.fr) (grants from the Conseil Régional d’Aquitaine n°20030304002FA and 20040305003FA, from the European Union FEDER n°2003227 and from Investissements d’Avenir ANR-10-EQPX-16-01). They also thank Géraldine Gourgues, Dr. Fabien Labroussaa and Angélique Alonso-Marrau for skilled technical assistance, Dr. Iason Tsarmpopoulos for providing biological material and advices and Pr. Alain Blanchard for revising the manuscript.

This work is part of the European MiniCell project « A model-driven approach to minimal cell engineering for medical therapy » selected by ANR, in the frame of the ERASynBio 2nd Joint Call for Transnational Research Projects (N° ANR-15-SYNB-0001-04). It has also been supported by the National Science Foundation [grant number IOS-1110151] and the European Union’s Horizon 2020 research and innovation program under grant agreement N°634942.

