## Supplementary Figure S1-S11 and Supplementary Table S1-S2 for "CReasPy-cloning: a method for simultaneous cloning and engineering of megabase-sized genomes in yeast using the CRISPR-Cas9 system"

**Authors and affiliation**:

Estelle Ruiz ^1,2^, Vincent Talenton^1,2^, Marie-Pierre Dubrana^1,2^, Gabrielle Guesdon^1,2^, Maria Lluch-Senar^3,4^, Franck Salin^5^, Pascal Sirand-Pugnet^1,2^, Yonathan Arfi^1,2^, Carole Lartigue^1,2*^

^1^ INRA, UMR 1332 de Biologie du Fruit et Pathologie, F-33140 Villenave d'Ornon, France

^2^ Univ. Bordeaux, UMR 1332 de Biologie du Fruit et Pathologie, F-33140 Villenave d'Ornon, France

^3^ EMBL/CRG Systems Biology Research Unit, Centre for Genomic Regulation (CRG), The Barcelona Institute of Science and Technology, Dr Aiguader 88, Barcelona 08003, Spain.

^4^ Universitat Pompeu Fabra (UPF), 08003 Barcelona, Spain.

^5^ BIOGECO, INRA, Univ. Bordeaux, 33610 Cestas, France

^*^**Corresponding author:**

INRA, Centre de Recherche de Bordeaux, UMR 1332 Biologie du Fruit et Pathologie, 71, avenue Edouard Bourlaux, CS 20032, 33882 Villenave d’Ornon Cedex, France.

**This file includes**

Supplementary Figures S1 to S11

Supplementary Tables S1 to S2

Puro-R

HIS3

ARS

CEN

Puro-R

HIS3

CEN

HR

HR

HR

HR

**Figure S1. Schematic view of the yeast elements recombination cassettes used for the CReasPy-cloning.** Top: “+ ARS” cassette – 3,115 bp; Bottom: “- ARS” cassette – 2,638 bp. Both cassette contains (i) two 60 bp recombination arms identical to the *M. pneumoniae* locus to edit (“HR”), (ii) a Histidine auxotrophic selection marker (“HIS3”) and (iii) a puromycine resistance marker (“Puro-R”)

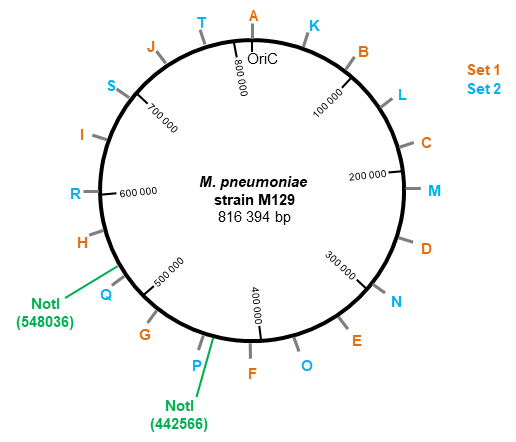

**Figure S2. Graphical map of *M. pneumoniae* M129 genome, displaying the location of the two sets of multiplex primers (set 1 and set 2).** Each set is composed of ten pairs of primers, allowing the amplification of 10 fragments, evenly distributed all around the genome and ranging from 100 to 1,000 bp (set 1) or from 125 to 1,025 bp (set 2), in 100 bp increments. The location of the two NotI restriction sites used for PFGE analysis is also displayed.

**Figure 2A**

**Figure 2B Top**

**Figure 2B Bottom**

**Figure 2C**

**Figure S3.** Full images used to produce the panels in Figure 2. The sections spliced together are highlighted with blue rectangles.

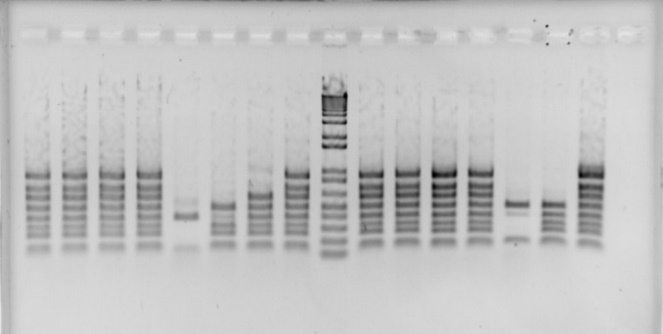

**+**

**-**

**M**

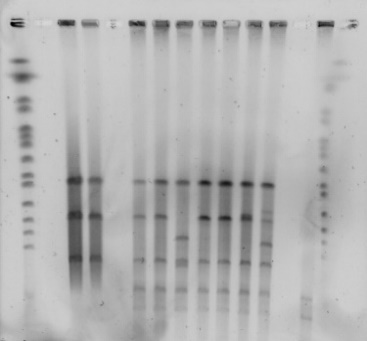

**+**

**M**

**+**

**A**

**B**

**C**

**D**

**8**

**9**

**13**

**10**

**17**

**21**

**33**

**34**

**36**

**37**

**39**

**40**

**44**

**46**

**8**

**9**

**13**

**34**

**36**

**39**

**40**

**M**

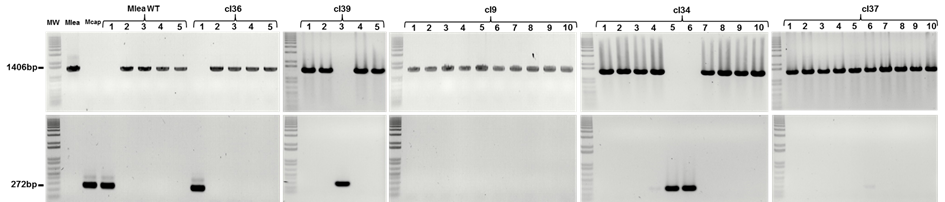

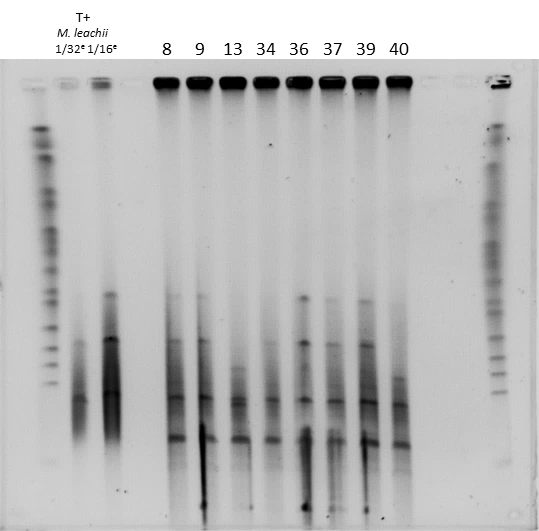

**36**

**37**

**40**

**39**

**M**

**13**

**34**

**8**

**9**

**M**

**+	+**

**46**

**M**

**M**

**M**

**1**

**2**

**3**

**4**

**5**

**6**

**7**

**8**

**9**

**10**

**11**

**12**

**13**

**14**

**15**

**16**

**17**

**18**

**19**

**20**

**21**

**22**

**23**

**24**

**25**

**26**

**27**

**28**

**29**

**30**

**31**

**32**

**33**

**34**

**35**

**36**

**38**

**37**

**39**

**40**

**41**

**42**

**43**

**44**

**45**

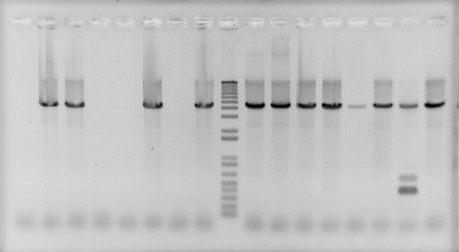

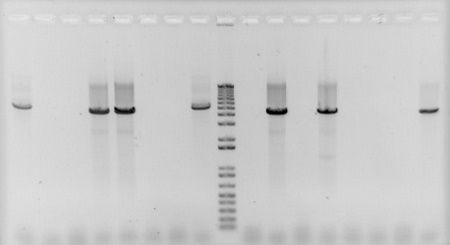

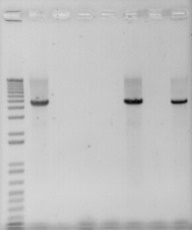

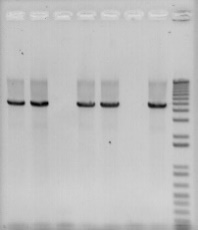

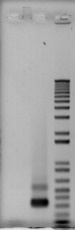

**M**

**M**

**-**

**+**

**Size (bp)**

**2000 -**

**1000 -**

**500 -**

**5000 -**

**Size (bp)**

**2000 -**

**1000 -**

**500 -**

**5000 -**

**300 -**

**100 -**

**Figure S4. Screening of VL6-48N yeast transformants carrying *Mycoplasma leachii* strain PG50 genome with deletion of MSB_0138 gene**. **(A)** Detection of *M. leachii* genome in yeast and of the expected deletion of the target gene by simplex PCR analysis with *M. leachii* specific primers designed inside the target gene. The expected sizes of the PCR products before and after replacement of the internal part of the MSB_0138 gene by the recombination template are 332 bp and 4,679 bp, respectively. “M”: DNA Ladder; “+”: *M. leachii* strain PG50 gDNA; “-”: negative control without DNA; **(B)** Screening for *M. leachii* genome completeness by multiplex PCR using a set of ten pairs of primers evenly distributed all around the *M. leachii* genome allowing the simultaneous amplification of ten fragments (~100 bp to 1,000 bp). “M”: DNA Ladder; “+”: *M. leachii* strain PG50 gDNA; “-”: negative control without DNA. **(C)** Validation of *M. leachii* genome integrity by XhoI digestion (three bands at 548 kbp, 121 kbp and 339 kbp are expected) and pulsed-field gel electrophoresis (PFGE). During the first PFGE (left panel), the yeast clone 37 was omitted. A second PFGE was performed (right panel). While the quality of the picture is poor and the 3-bands pattern hardly seen for some clones, the clone 37 showed the expected pattern and was selected together with yeast clones 9, 34, 36 and 39 for the following step (genome transplantation). “M”: PFGE marker; “+”: *M. leachii* strain PG50. **(D)** Validation of mutants produced by genome transplantation of CReasPy-cloned *M. leachii* genomes, by simplex PCR using *M. leachii* specific primers. *M. leachii* transplants are expected to show a 1,406 bp amplicon, while spontaneously resistant recipient cells would show no amplification. “M”: DNA Ladder; “+”: *M. leachii* strain PG50 gDNA.

**A**

**B**

**16.1**

**16.2**

**16.3**

**16.4**

**16.5**

**M**

**+**

**16.6**

**16.7**

**16.10**

**M**

**-**

**+**

**Size (bp)**

**100 -**

**300 -**

**500 -**

**1000 -**

**1517 -**

**16.10**

**16.6**

**16.7**

**16.8**

**16.9**

**-**

**+**

**M**

**-**

**+**

**M**

**16.1**

**16.2**

**16.3**

**16.4**

**16.5**

**Size (bp)**

**2000 -**

**1000 -**

**500 -**

**5000 -**

**Size (bp)**

**2000 -**

**1000 -**

**500 -**

**5000 -**

**Size (kb)**

**M**

**16.2**

**16.6**

**16.7**

**+**

**M**

**M**

**16.2**

**16.6**

**16.7**

**+**

**225 -**

**295 -**

**375 -**

**450 -**

**553 -**

**680 -**

**745 -**

**815 -**

**945 -**

**1640 -**

**1900 -**

**M**

**915 -**

**Size (kb)**

**225 -**

**375 -**

**450 -**

**553 -**

**680 -**

**745 -**

**815 -**

**945 -**

**1100 -**

**1900 -**

**C**

**Figure S5. Screening of yeast transformants carrying *Mycoplasma mycoides* subsp. m*ycoides* strain Afadé genome deleted for TS60_0301-0299 operon. (A)** Detection of *M. mycoides* subsp*. mycoides* (*Mmm*) genome and of the correct deletion of the target gene by simplex PCR analysis using specific primers designed on either side of the target gene. The expected size after substitution by the recombination template is 4,588 bp, whereas 4,022 bp correspond to the size of the wild-type TS60_0301-0299 operon. **(B)** Screening for *Mmm* genome completeness by multiplex PCR using a set of nine pairs of primers evenly distributed all around the *Mmm* genome, allowing the simultaneous amplification of nine fragments ranging from 100 to 1,000 bp in 100 bp increments. “M”: DNA Ladder; “+”: *M. mycoides* subsp. *mycoides* strain Afadé gDNA; “-”: negative control without DNA. **(C)** The size of the *Mmm* genome cloned in yeast is assessed by enzymatic restriction and Pulsed Field Gel Electrophoresis (PFGE). Left: Digestion by the restriction enzyme BssHII should yield one linear DNA fragments of ~1,2 Mbp. Right: Digestion by the restriction enzyme XhoI should yield three linear DNA fragments of 950 kb, 161 kb and 76 kb. “M”: PFGE DNA ladder; “+”: *M. mycoides* subsp*. mycoides*

**Figure 4A**

**Figure 4B**

**Figure 4C**

**Figure 4D**

**Figure 4E**

**Figure S6.** Full images used to produce the panels in Figure 4. The sections spliced together are highlighted with blue rectangles

+ *In vitro* transcribed gRNA

+ Recombinant Cas9

Cells embedded

in agarose

+ Proteinase K

+ Detergent

X

X

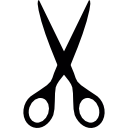

+ β-agarase

+ heat

Linearized

chromosome

Yeast

elements

Removal of cellular

Envelope and protein digestion

Cleavage of the

circular chromosome

Co-transformation

**Figure S7. *In vitro* CReasPy-cloning**. Cells embedded in agarose plugs are digested to obtain naked DNA embedded in agarose plugs. Then, the DNA is incubated *in vitro* in presence of Cas9 and gRNA, prior to yeast transformation. The digested DNA and an appropriate recombination template are co-transformed in yeast, resulting in the repair of the *M.* *pneumoniae* genome by the highly efficient yeast homologous recombination system.

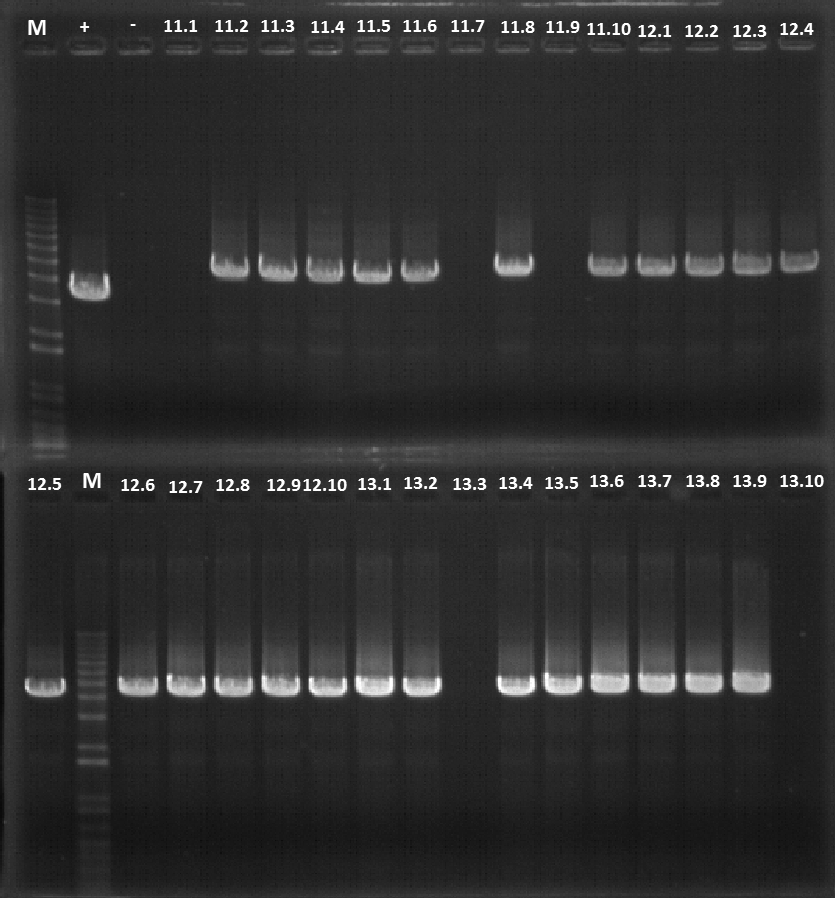

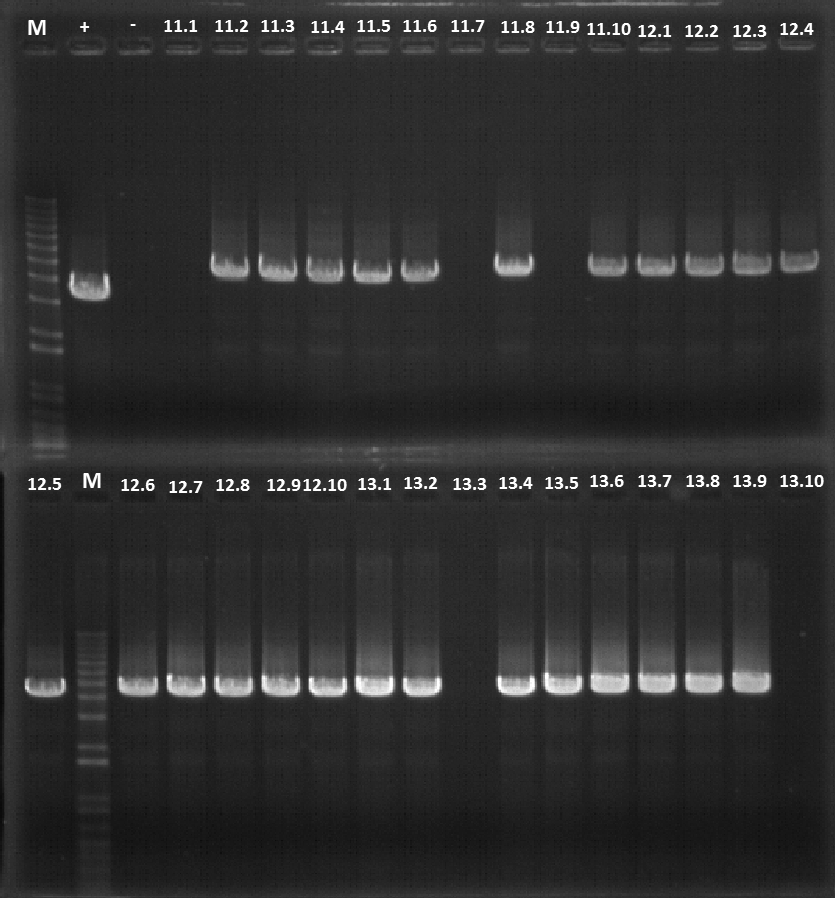

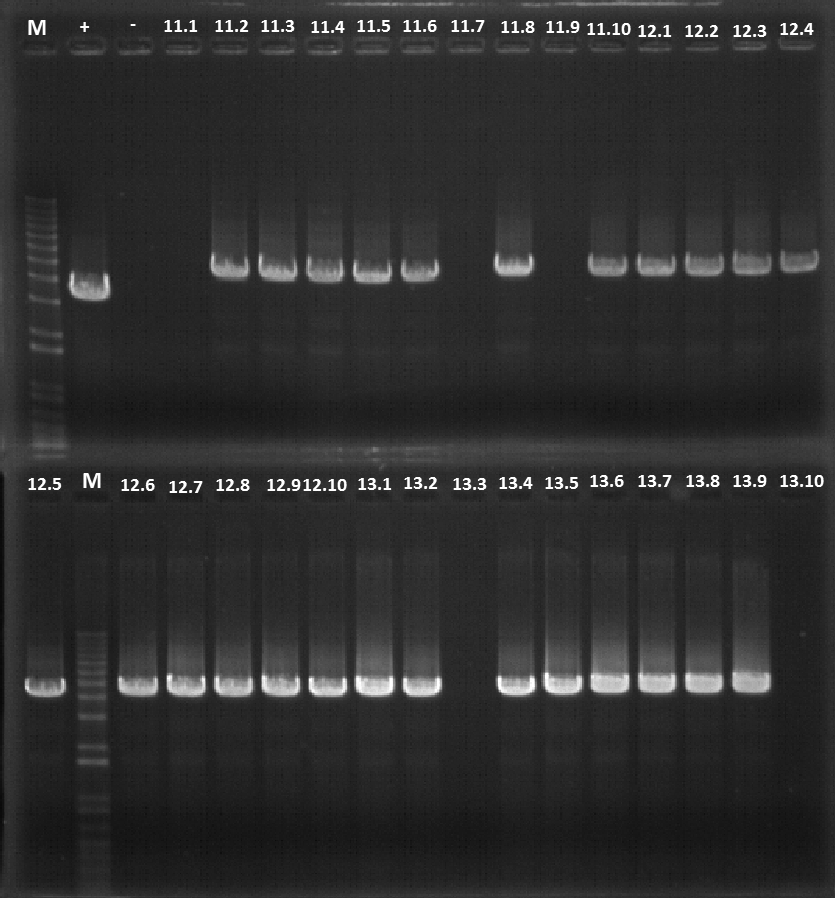

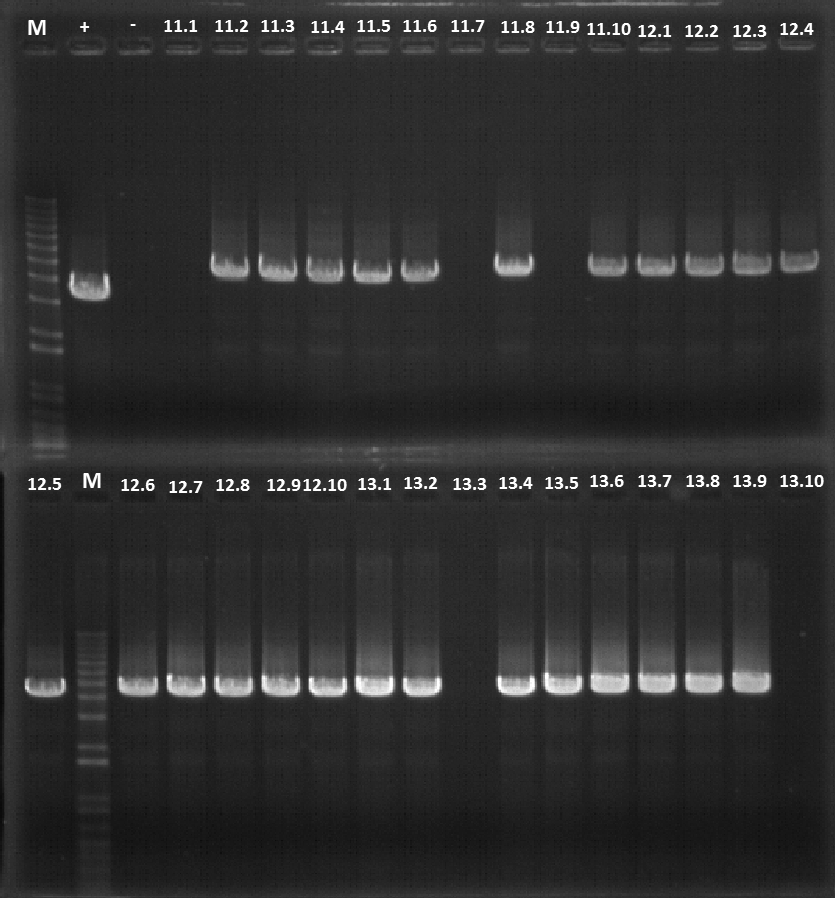

**12.1**

**12.2**

**12.3**

**12.4**

**M**

**+**

**-**

**12.5**

**12.6**

**12.7**

**12.8**

**12.9**

**12.10**

**Size (bp)**

**2000 -**

**1000 -**

**500 -**

**5000 -**

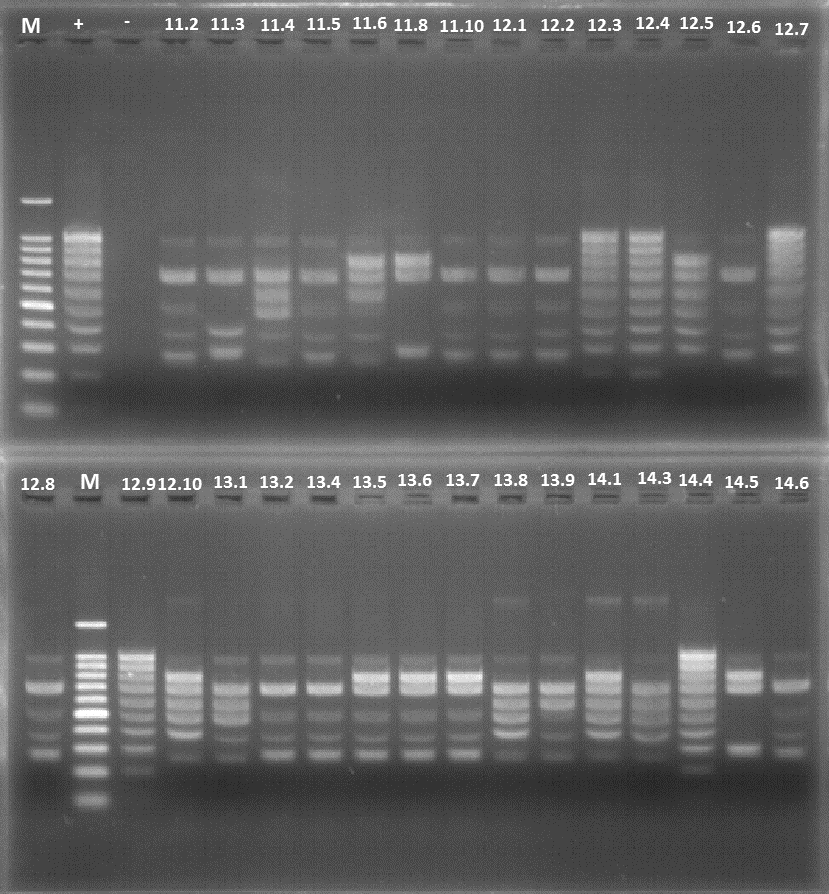

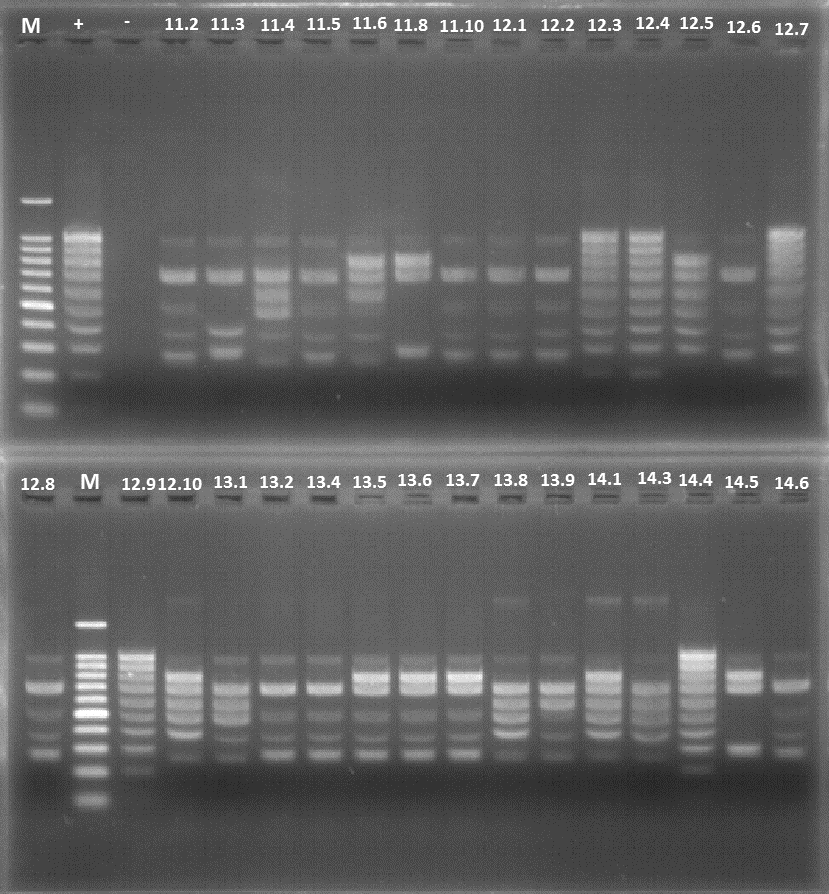

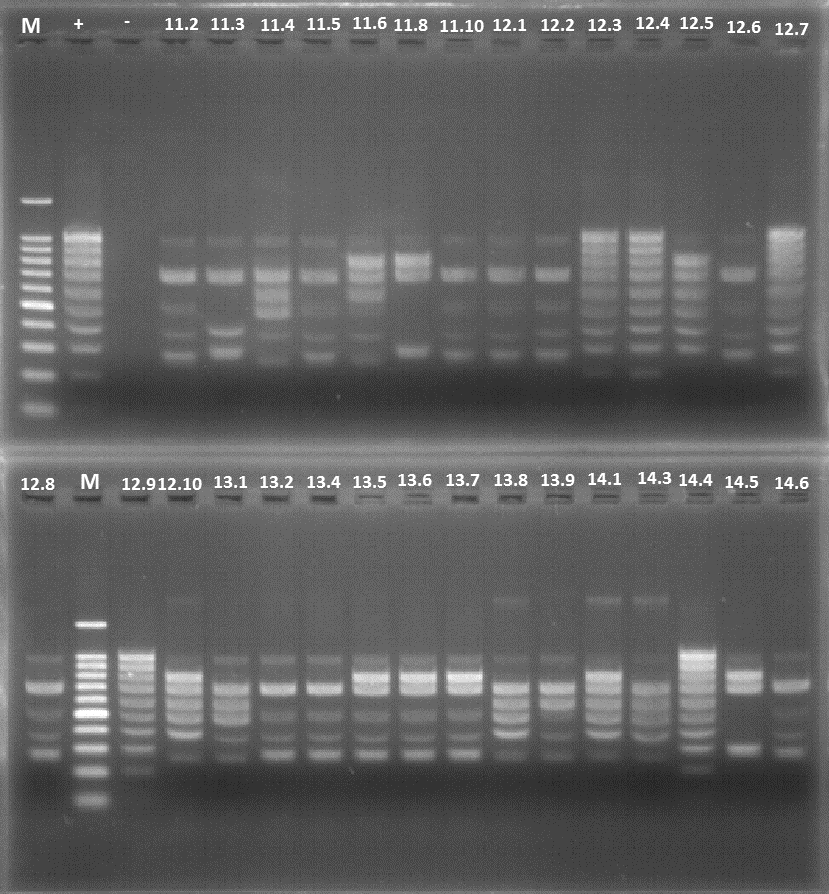

**12.1**

**12.2**

**12.3**

**12.4**

**M**

**+**

**-**

**12.5**

**12.6**

**12.7**

**12.8**

**12.9**

**12.10**

**Size (bp)**

**M**

**1000 -**

**500 -**

**300 -**

**100 -**

**700 -**

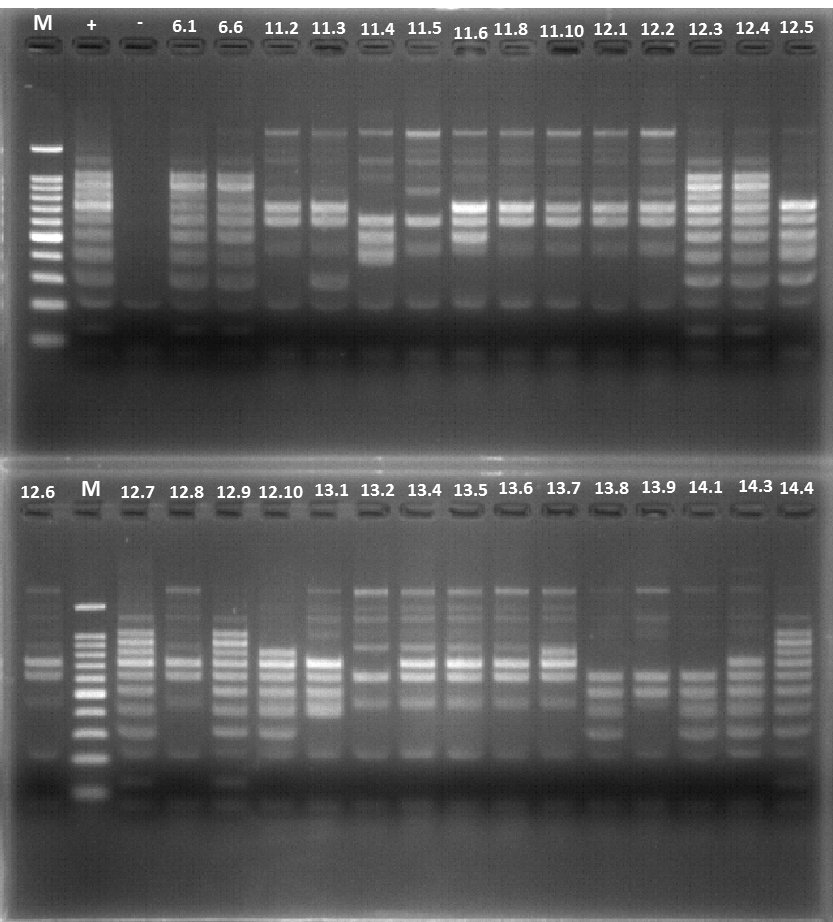

**12.1**

**12.2**

**12.3**

**12.4**

**M**

**+**

**-**

**12.5**

**12.6**

**M**

**12.7**

**12.9**

**12.10**

**Size (bp)**

**1000 -**

**500 -**

**300 -**

**100 -**

**700 -**

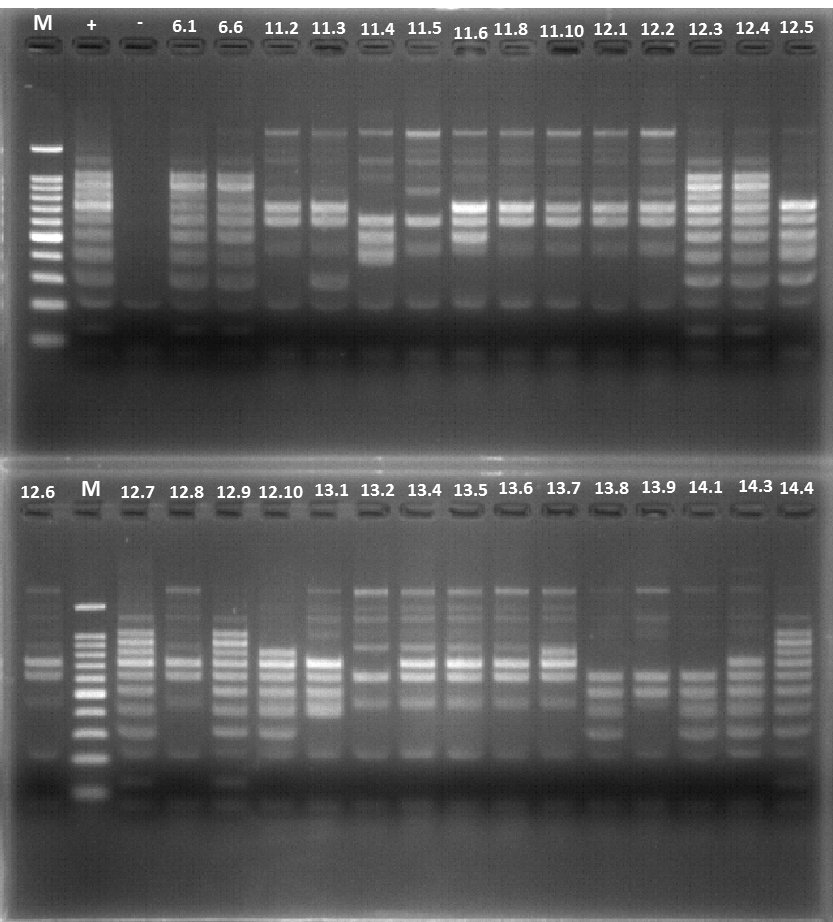

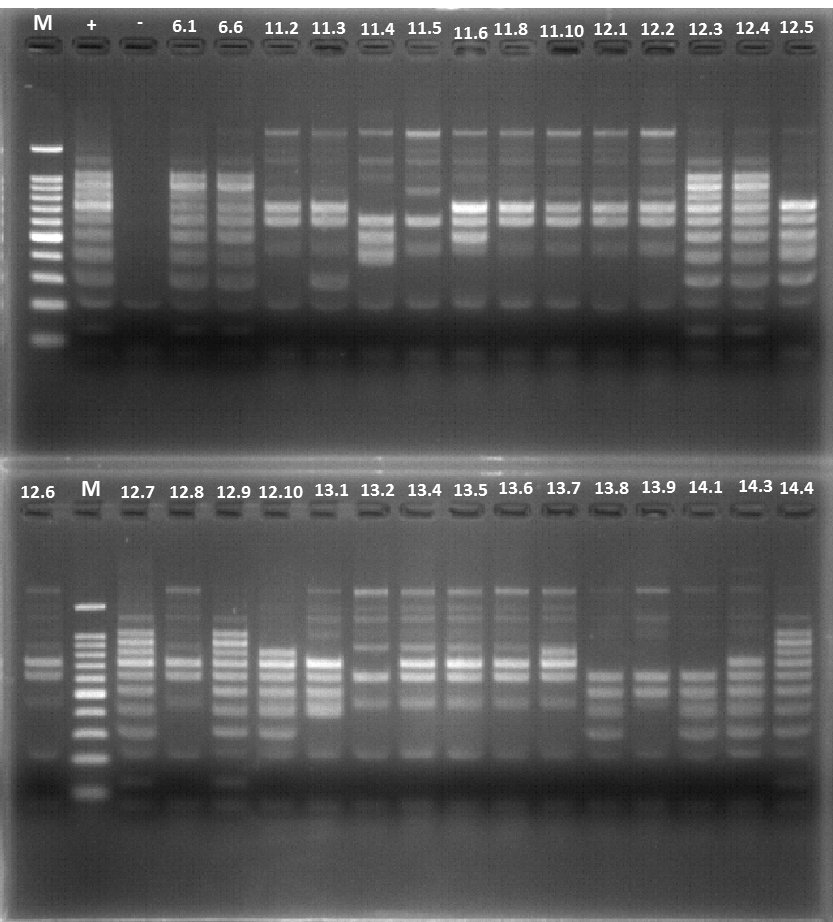

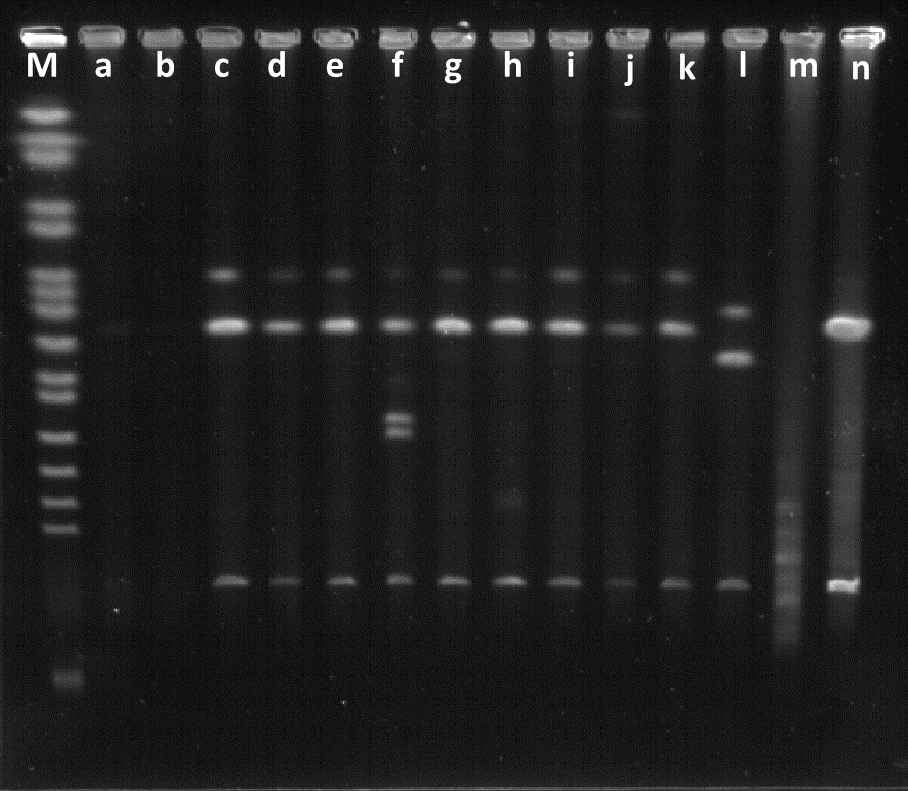

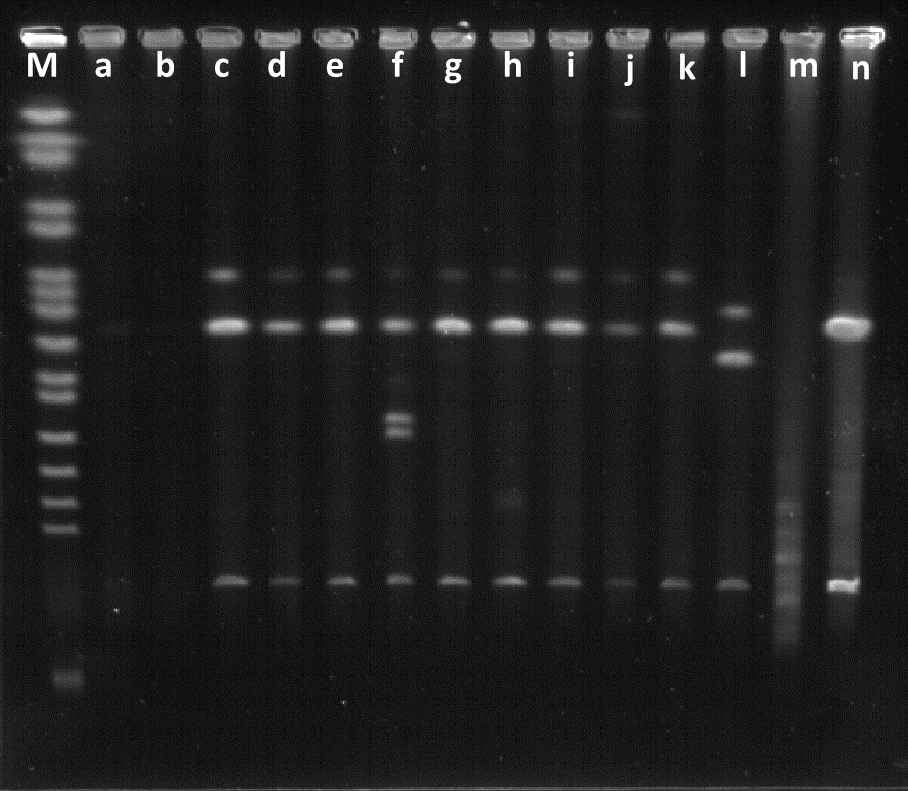

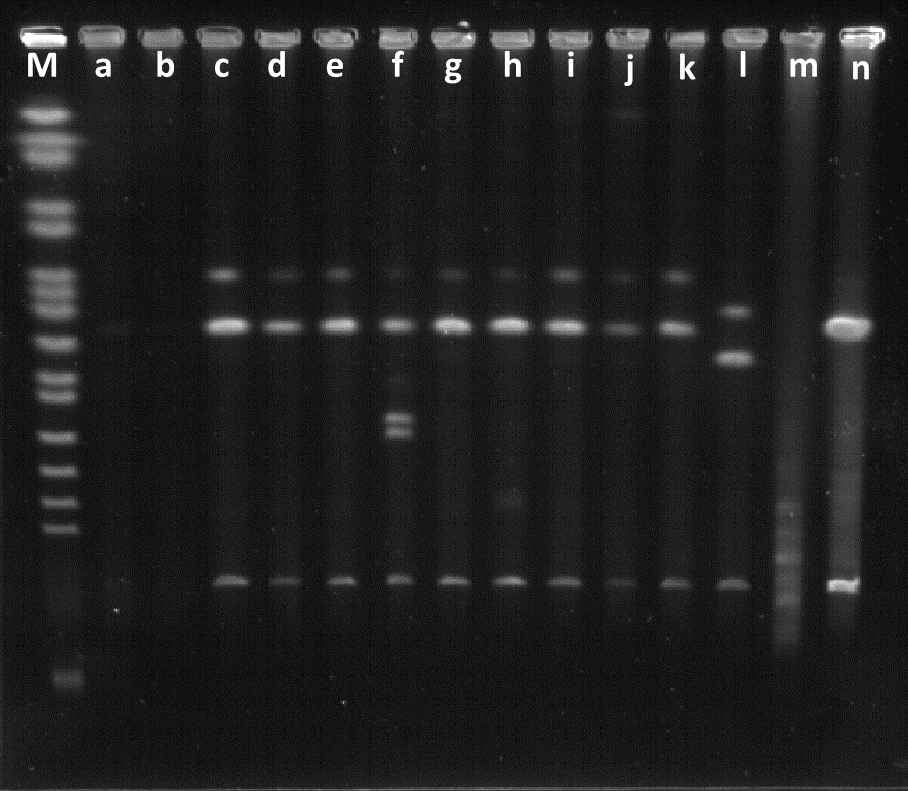

**Sc**

**12.3**

**12.4**

**12.7**

**M**

**Size (kbp)**

**225 -**

**295 -**

**375 -**

**450 -**

**553 -**

**610 -**

**680 -**

**745 -**

**815 -**

**12.9**

**+**

**915 -**

**945 -**

**A**

**B**

**C**

**12.8**

**Figure S8**. **Screening of yeast transformants generated by *in vitro* CReasPy-cloning of *M. pneumoniae* M129 genome and deletion of MPN400**. The images used to produce panels A-C are spliced from multiple gels in order to display the same 10 yeast clones. Original images are available in Supplementary Figure S9. (A) The presence of the *M. pneumoniae* genome in the yeast and the desired target replacement were checked by simplex PCR analysis. Using *M. pneumoniae* specific primers flanking the target gene, an amplicon of 4,841 bp was expected in properly edited clones, whereas a fragment of 3,480 bp was expected for wild types clones. (B-C) The completeness of the *M. pneumoniae* genome cloned in yeast was assessed by multiplex PCR using two sets of primers (Top: set 1, Bottom: set 2). Each set was comprised of ten pairs of primers evenly distributed around the genome of *M. pneumoniae*, allowing the simultaneous amplification of ten fragments ranging from 100 to 1,000 bp (set 1) and from 125 to 1,025 bp (set 2) in 100 bp increments. Clones carrying *M. pneumoniae* genomes without major rearrangement displayed a ten bands profile identical to the one obtained in the positive control. “M”: DNA Ladder; “+”: *M. pneumoniae* M129 gDNA; “-”: negative control without DNA. (C) The size of the *M. pneumoniae* genome cloned in yeast was assessed by enzymatic restriction and Pulsed Field Gel Electrophoresis (PFGE). Digestion by the restriction enzyme NotI-HF^®^ should yield two linear DNA fragments of 711 kb and 105 kb. “M”: PFGE DNA ladder; “Sc”: *S. cerevisiae* VL6-48N; “+”: *M. pneumoniae* M129

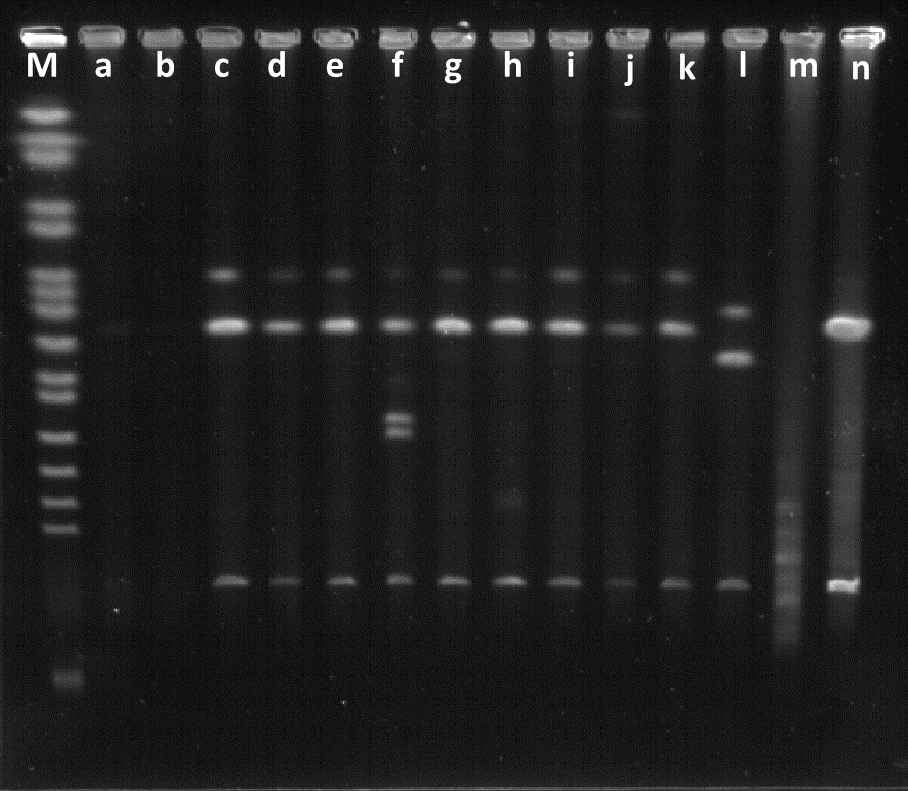

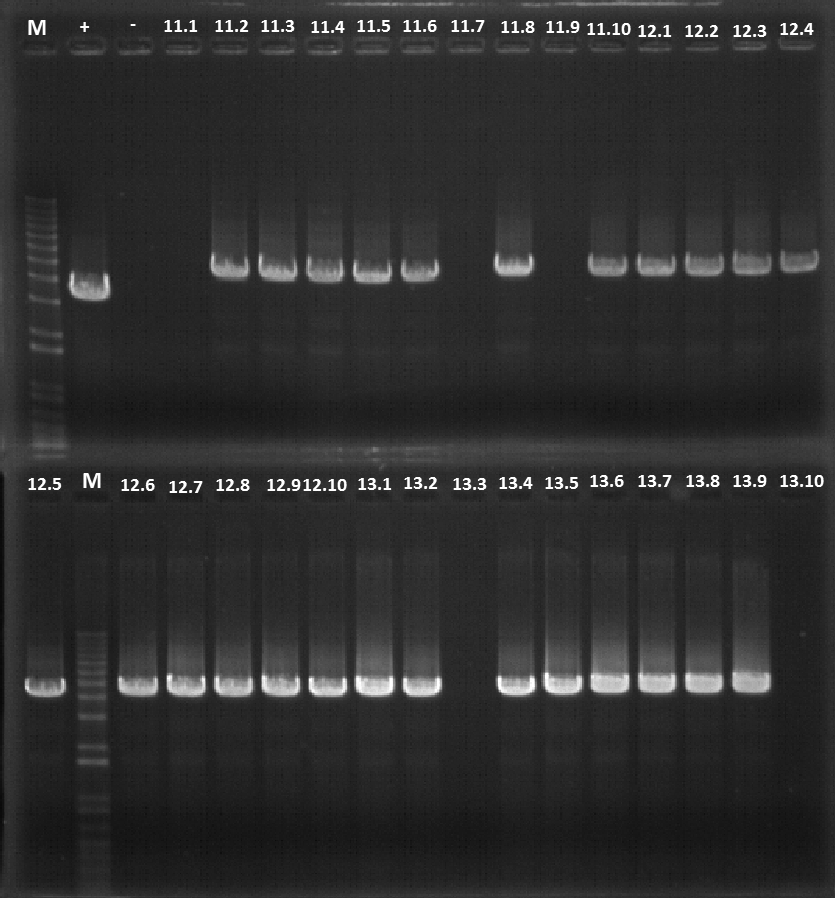

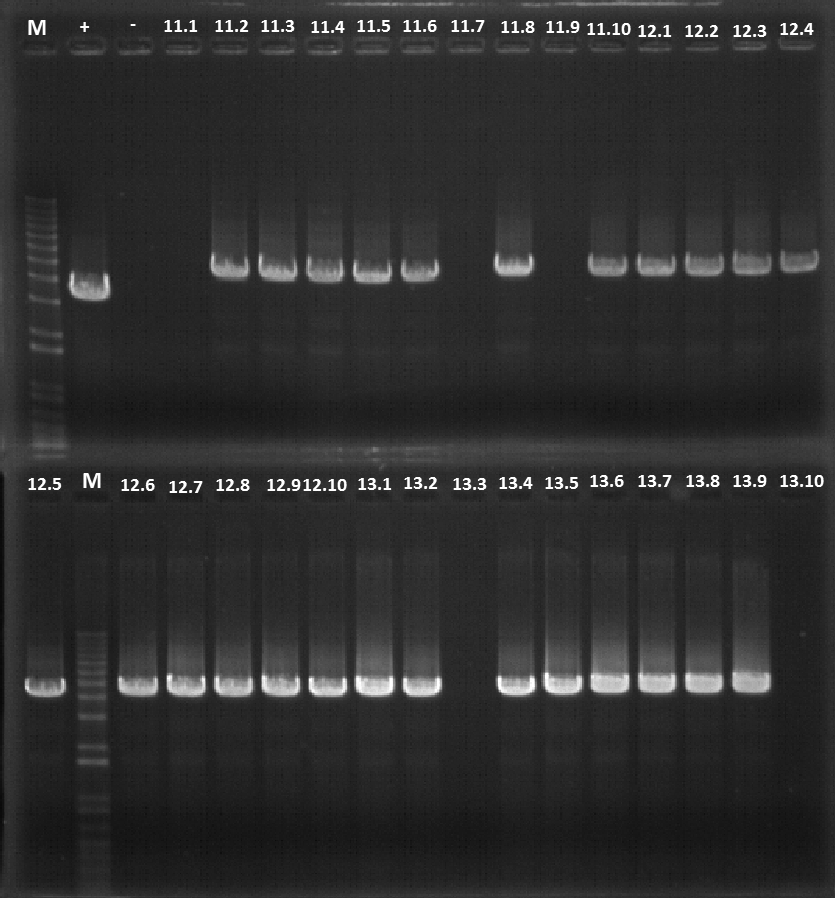

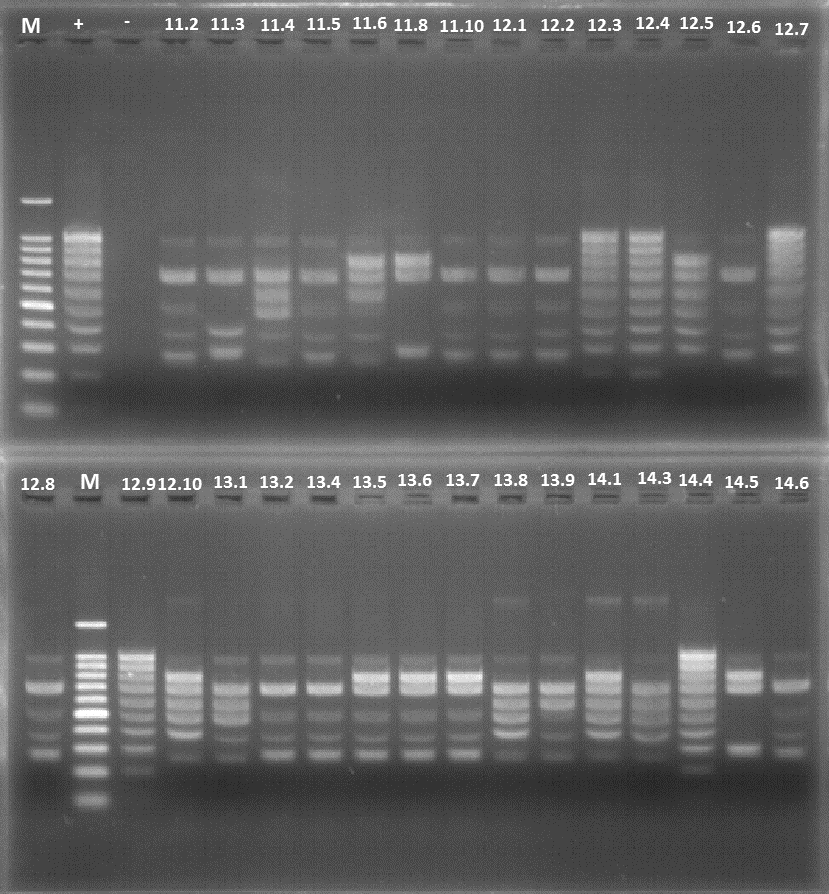

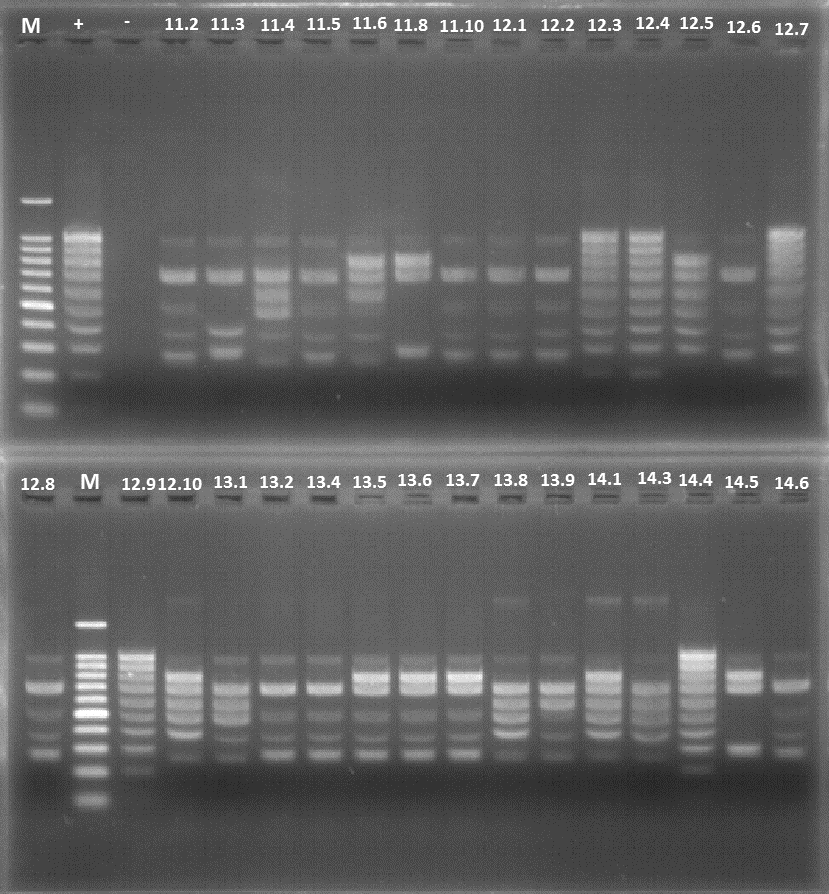

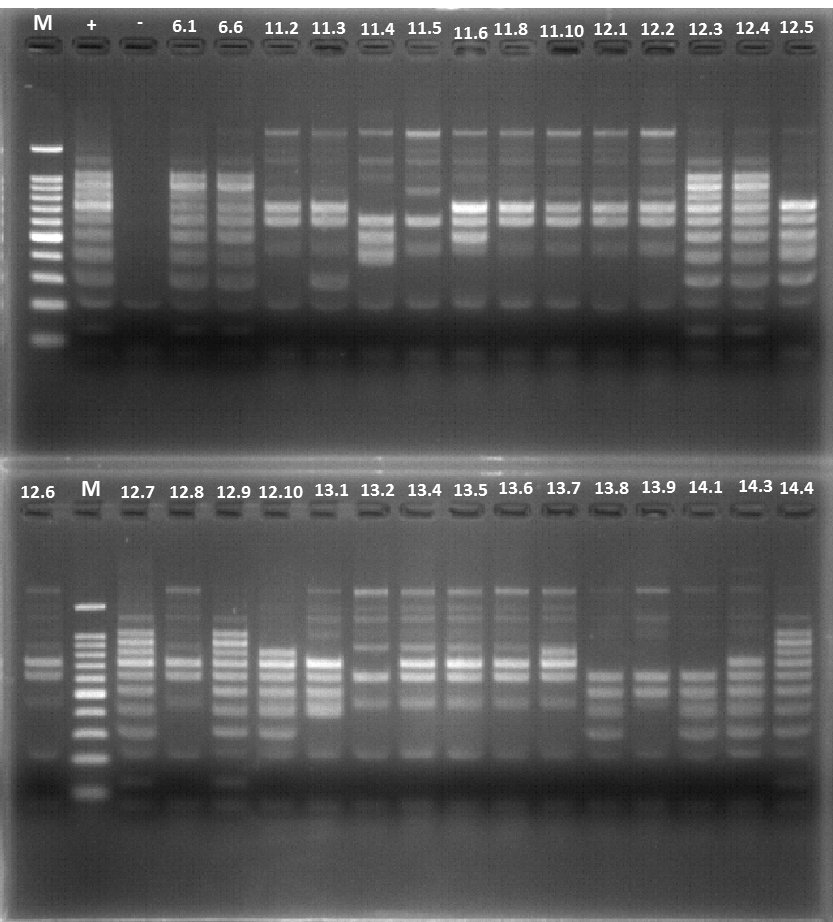

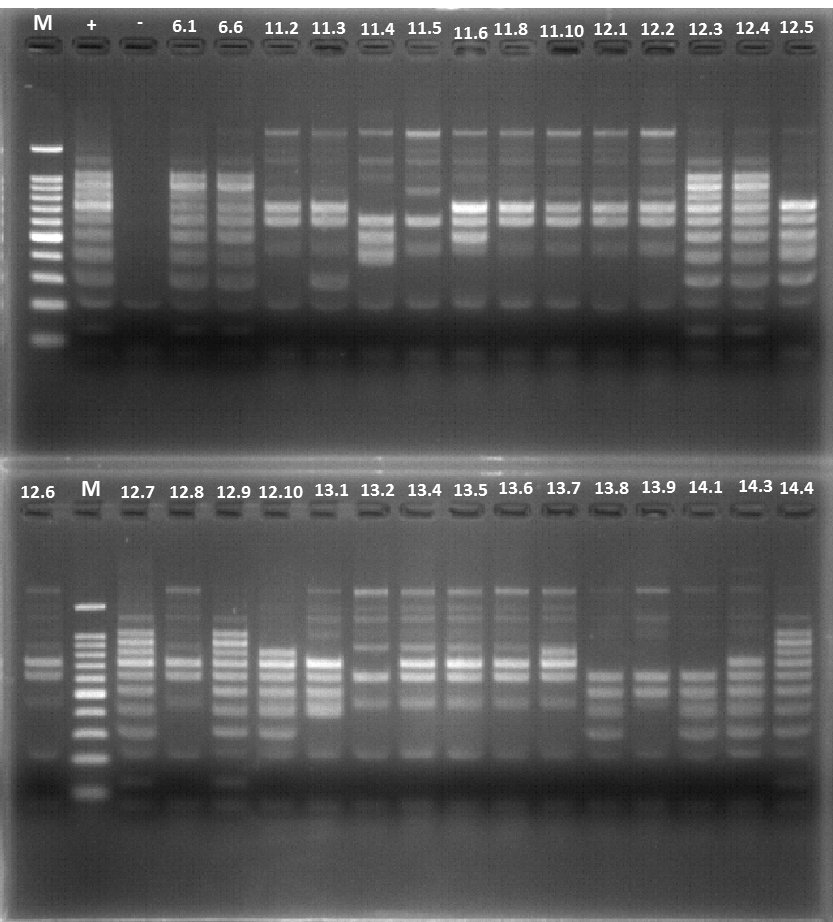

**Figure S8A**

**Figure S8B Bottom**

**Figure S8B Top**

**Figure S8C**

**Figure S9.** Full images used to produce the panels in Figure S8. The sections spliced together are highlighted with blue rectangles

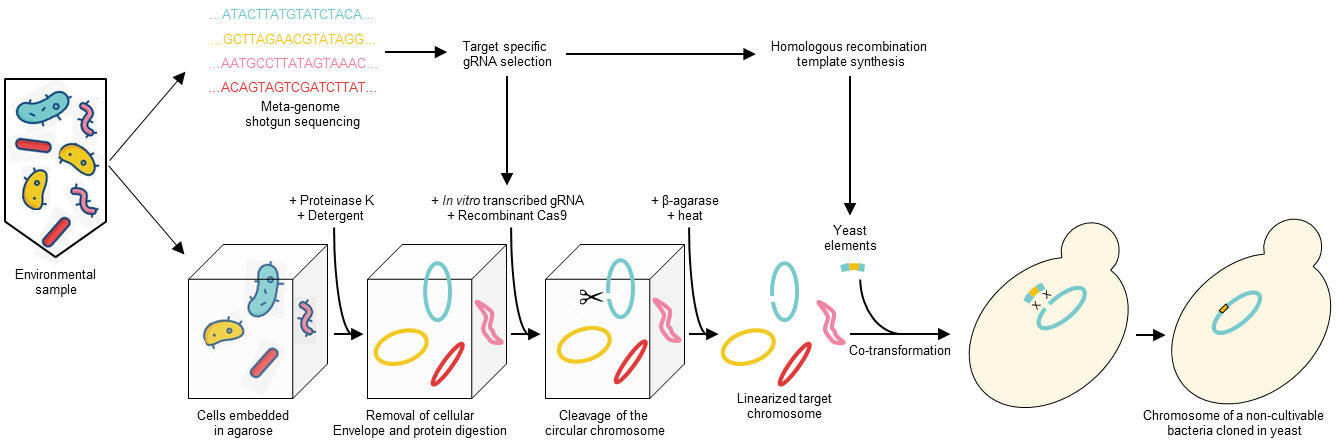

**Figure S10.** Proposed application of CReasPy-cloning to capture the genome of uncultivable micro-organisms. An environmental sample containing a mixture of cultivable and uncultivable species is subdivided in two parts: one is analyzed by shotgun sequencing in order to collect metagenomics data; the other is encased in agarose plugs in order to isolate the intact chromosomes of the micro-organisms. The metagenomics data are used to select a suitable target in a genome of interest, and to produce the appropriate gRNA and recombination cassette. Using *in vitro* CReasPy-cloning, the target chromosome is linearized and subsequently transformed in yeast. After repair, the genome of the uncultivable organism is now carried in yeast, and can be fully sequenced and modified.

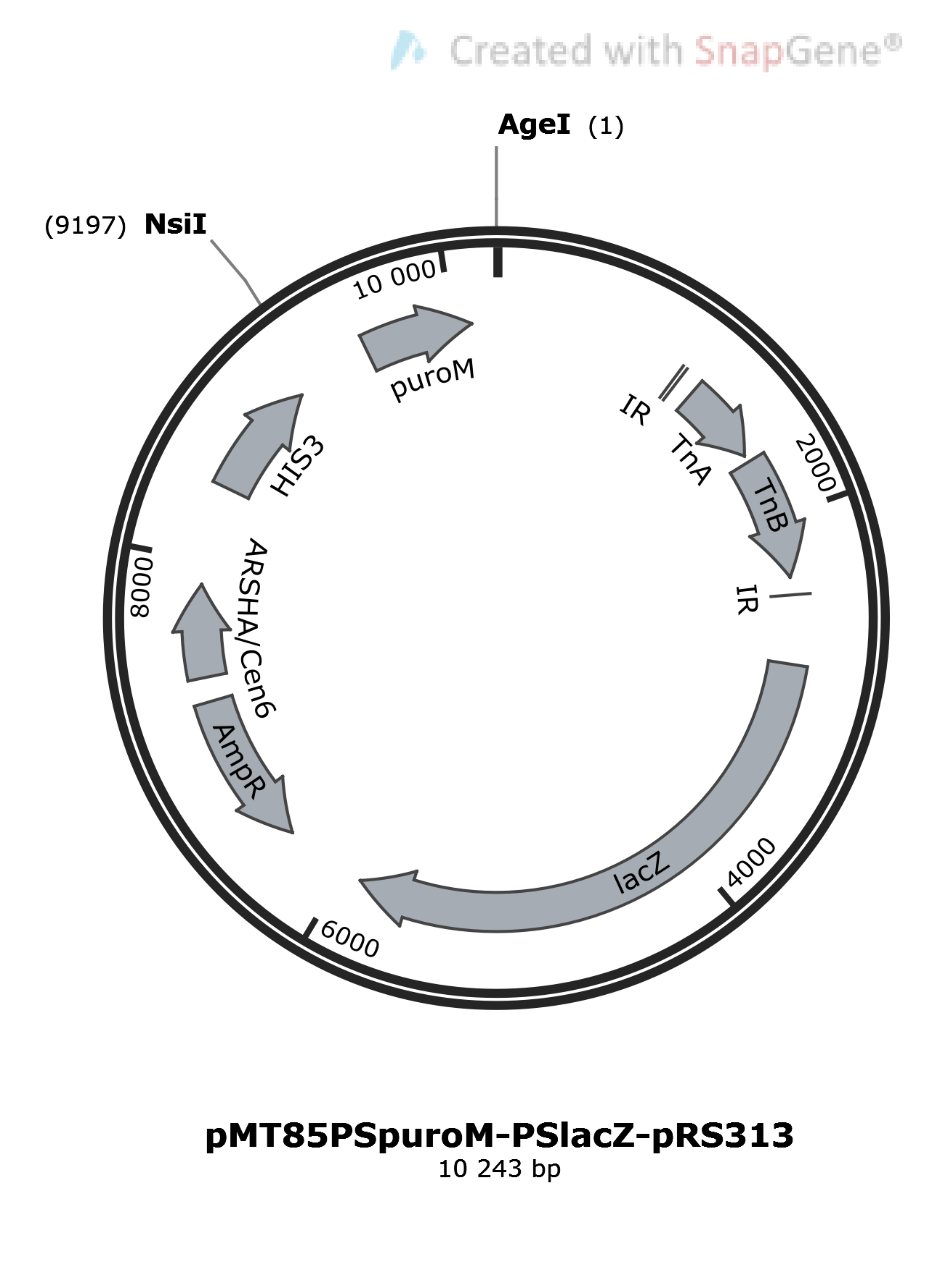

**10,243 bp**

**Figure S11.** Map of the plasmid pMT85PSpuroM-PSlacZ-pRS313. The puromycin cassette was amplified from the pMiniTn4001PSpuro (Algire et al, 2009) using primers that contain NsiI or AgeI restriction sites (PSpuroNsiI, 5’-CATGATGCATGTGAACAAGAAAACAGTGAAGC; PSpuroAgeI, 5’-CATGACCGGTCATGCCTGCAGTTCAGATCC). The ligation of the puromycin cassette with the pMT85PStetM-PSlacZ-pRS313 (Genbank accession number KX011460, Labroussaa et al, 2016) both digested with AgeI-HF® and NsiI-HF® resulted in the pMT85PSpuroM-PSlacZ-pRS313 construction ’10,243 bp).

**Table S1.** Assessment of the number of genetic elements that can be co-transformed into yeast during CReasPy-cloning.

| **Yeast strain** | **Elements co-transformed**  **into yeast** | **Simplex PCR screening^a^:**  **Positive clones /**  **Analyzed clones** |
| --- | --- | --- |
| *S. cerevisiae* VL6-48N | *M. pneumoniae* chromosome  Recombination template  pgRNA  pCas9 | 0/7 |
| *S. cerevisiae* VL6-48N-pCas9 | *M. pneumoniae* chromosome  Recombination template  pgRNA | 0/8 |
| *S. cerevisiae* VL6-48N-pCas9-pgRNA | *M. pneumoniae* chromosome  Recombination template | 9/10 |

^a^ Different CReasPy-cloning experiments were performed before, during which the target chromosome and the recombination template were co-transformed with either the pCas9 and pgRNA, or the pgRNA. The elements that are not co-transformed were introduced in the yeast beforehand. Yeast transformants were screened by simplex PCR to validate the proper deletion of MPN372 by the cassette. The number of positive clones is given for each experiment.

**Table S2.** Comparison of the CReasPy-cloning efficiency depending on the utilization of a yeast element cassette containing or not an ARS sequence.

|  | **Positive clones / Analyzed clones** | | |
| --- | --- | --- | --- |
| **Recombination template** | **Targeted locus** | **Simplex PCR** | **Multiplex PCR** |
| + ARS | MPN372 | 10/20 | 2/20 |
|  | MPN142 | 1/20 | 1/1 |
|  | MPN142-143 | 4/20 | 2/4 |
|  | MPN400 | 10/20 | 4/10 |
|  | MPN398-400 | 8/20 | 2/8 |
| - ARS | MPN372 | 20/20 | 16/20 |
|  | MPN142 | 19/20 | 15/19 |
|  | MPN142-143 | 16/20 | 11/16 |
|  | MPN400 | 20/20 | 10/20 |
|  | MPN398-400 | 20/20 | 14/20 |
